# Characterization and Prediction of ISRE Binding Patterns Across Cell Types Under Type I Interferon Stimulation

**DOI:** 10.1101/2020.09.08.287581

**Authors:** Sivan Leviyang

## Abstract

Stimulation of cells by type I interferons (IFN) leads to the differential expression of 100s of genes known as interferon stimulated genes, ISGs. The collection of ISGs differentially expressed under IFN stimulation, referred to as the IFN signature, varies across cell types. Non-canonical IFN signaling has been clearly associated with variation in IFN signature across cell types, but the existence of variation in canonical signaling and its impact on IFN signatures is less clear. The canonical IFN signaling pathway involves binding of the transcription factor ISGF3 to IFN-stimulated response elements, ISREs. We examined ISRE binding patterns under IFN stimulation across six cell types using existing ChIPseq datasets available on the GEO and ENCODE databases. We find that ISRE binding is cell specific, particularly for ISREs distal to transcription start sites, potentially associated with enhancer elements, while ISRE binding in promoter regions is more conserved. Given variation of ISRE binding across cell types, we investigated associations between the cell type, homeostatic state and ISRE binding patterns. Taking a machine learning approach and using existing ATACseq and ChIPseq datasets available on GEO and ENCODE, we show that the epigenetic state of an ISRE locus at homeostasis and the DNA sequence of the ISRE locus are predictive of the ISRE’s binding under IFN stimulation in a cell type, specific manner, particularly for ISRE distal to transcription start sites.

## 1 Introduction

Stimulating cells with type I interferons (IFN), in particular IFNa and IFNb, activates signaling pathways that lead to the upregulation of a collection of genes known as interferon stimulated genes, ISGs. ISGs play a diverse and essential role in the innate immune response, with some acting directly as antiviral effectors while others regulate the innate and adaptive responses [68]. Cell types vary in their ISGs, presumably reflecting the need for different functional responses to infection. For example, IFNb stimulation of monocytes and B cells leads to 100s of ISGs in both cell types, but only 40% of the ISGs are shared in the two cell types, and the difference in ISGs associates with cell death in monocytes and proliferation in B cells [76]. Neuronal cell types differ in their upregulation of the antiviral genes IFI27, IRG1 and RSAD2 under IFNb stimulation, leading to variation in viral restriction [13]. Stem cells constitutively upregulate many genes that are ISGs in other cell types, possibly to allow for viral restriction while avoiding IFN mediated antiproliferative effects [84]. IFN stimulation of cardiac myocytes weakly induces some ISGs that are potently induced in cardiac fibroblasts, possibly reflecting functional restrictions on cardiac monocytes which are not replenished [88].

Despite the profound differences in ISGs across cell types, our understanding of the cellular factors and pathways that determine a cell type’s ISGs, referred to as the cell type’s IFN signature [4], is incomplete. The canonical IFN signaling pathway involves activation of STAT1 and STAT2 and the formation of the IGSF3 trimer, composed of STAT1, STAT2, and IRF9. ISGF3 binds to genomic loci called IFN-stimulated response elements, ISREs, and regulates the transcription of ISGs [20]. However, over the past two decades an array of non-canonical interferon pathways and regulators have been discovered [38]. In particular, IFN signaling can activate STATs other than STAT1 and STAT2, pathways other than JAK-STAT are activated by IFN stimulation or are involved in crosstalk with IFN signaling, and the STAT2:IRF9 dimer and other molecules other than ISGF3 can bind to ISREs [60, 83, 57, 25, 48, 61].

IFN signaling through non-canonical pathways has been clearly associated with variation in IFN signature. IFN signaling in immune cells activates STAT3 and STAT5, leading to profound differences in ISG signatures, e.g. [76, 35, 73, 82, 29, 36]. The balance between a type I IFN signature (induced by IFNa or IFNb) and type II IFN signature (induced by IFNg) is affected by cross talk between JAK-STAT and several pathways, including NFkB, e.g. [86, 54, 83, 37]. In work analyzing the different regulatory pathways influencing ISG expression, Mostafavi et al. grouped mouse ISGs into 5 modules and identified different transcription factors correlated with ISG expression levels in each module [52]. One module had ISGs particularly enriched in ISRE motifs and associated with ISGF3 component expression levels, but in other modules ISGs were not enriched for ISRE and were associated with transcription factors other than ISGF3.

However, while it is clear that IFN signaling through non-canonical pathways leads to variation in the IFN signature, the degree of variability in the IFN signature associated with canonical IFN signaling, mediated by binding of ISGF3 and other transcription factors to ISREs, is not clear. In an array of studies over the last decade, transcription factor binding has been shown to associate with epigenetic factors that vary between cell types [33, 71, 41, 22, 64, 53, 4, 72], but the extent to which this is the case for ISRE binding is unclear.

In this work, we took a step upstream of the IFN signature and considered the ISRE signature. By ISRE signature, we mean the collection of ISREs bound by ISGF3 and other transcription factors under IFN stimulation. We considered the ISRE signature across six cell types: mouse BMDM, fibroblast, and B cells, and human HeLa, K562, and THP1 cells. First, we characterized the ISRE signature across the six cell types, investigating the distribution of bound ISREs within genes and the overlap of bound ISREs across cell types. Second, we established an association between ISRE signatures and IFN signatures by correlating the presence of bound ISRE in gene regulatory regions with the gene’s status as an ISG. Third, we characterized the epigenetic state of the cell types at homeostasis and, through a machine learning approach, quantified the capacity of the homeostatic, epigenetic state along with genomic sequence specificities to predict ISRE binding under IFN stimulation.

Previous authors have investigated ISRE binding on a genomic scale. Seminal work by Hartman et al. and Robertson et al. identified STAT1 and STAT2 binding across the genome in HeLa cells, characterized ISRE motifs that were preferentially bound, and described overlap between STAT1 and STAT2 binding sites [32, 65]. Testoni et al. characterized STAT2 binding at homeostasis and under IFNa stimulation over a subset of 113 ISG promoters in hepatocytes. They found that STAT2 binding at homeostasis was associated with differential expression under IFN stimulation, but that STAT2 binding could act as a repressor [74]. Several groups have compared the effects of ISGF3 binding and STAT2:IRF9 dimer binding to ISREs [25, 7, 6, 61]. However, none of these works characterized the ISRE signature across the genome or across cell types.

The importance of epigenetic state in shaping the IFN response has been demonstrated in several contexts. Fang et al showed that levels of H3K9me2 modifications in promoters correlate inversely with ISG expression [24]. In the already mentioned work, Testoni et al. found an association between H3K4me1 and H3K27me3 markings at homeostasis and up and downregulation, respectively, of expression under IFN stimulation. Studies by Au-Yeung and Horvath, and Kadota and Nagata have shown that presence and absence of different histone types in promoters of ISGs regulate expression under IFN stimulation [1, 39]. Several studies have shown that external stimulus, for example in the form of LPS or IFN, leads to large scale epigenetic changes that regulate ISGs [52, 57, 40]. However, none of these works quantified the capacity of the cellular, epigenetic state at homeostasis to predict ISRE binding under IFN stimulation.

Our analysis provides several novel insights into the IFN response. Canonically, ISG regulation has been associated with ISRE binding in promoters. Our results support a role for bound ISRE distal to transcription start sites (TSS), presumably associated with enhancers. We find that most ISGs have bound ISREs in enhancers, that variation in the ISRE signature across cell types is predominantly associated with variation in enhancers, and that bound ISREs in enhancers associate with differential expression of ISGs, although more weakly than bound ISRE in promoters. We show that ISRE binding under IFN stimulation is predicted at relatively high accuracy by the homeostatic epigenetic state and sequence specificities of the ISRE, suggesting that factors controlling the homeostatic state of a cell play a substantial role in shaping the ISRE signature and, in turn, the IFN signature.

## 2 Materials and Methods

We considered the following cell types: mouse bone marrow derived macrophages (BMDM), embryonic fibroblasts (fibroblasts), and splenic B (B) cells, and human HeLa-S3 (HeLa), K562, and THP1 cells. All datasets were downloaded from the GEO or ENCODE databases. Tables 1, 2, and 3 provide references, and further details for ATACseq and histone modification ChIPseq datasets, STAT1, STAT2 and IRF9 ChIPseq datasets, and transcription datasets, respectively. See Supplementary Information for GEO and ENCODE accessions and number of replicates.

**Table 1:** Homeostatic, ATACseq and Histone Modification Datasets. For each cell type, we downloaded an ATACseq dataset and ChIPseq datasets for the H3K4ME1, H3K4ME3 and H3K27AC modifications. All datasets were collected under homeostatic conditions. See Supplementary Information for accessions and number of replicates.

| cell type | assay | reference |
| --- | --- | --- |
| BMDM | ATACseq | Czimmerer et al. [18] |
|  | H3K4ME1 | Ostuni et al. [56] |
|  | H3K4ME3 | Fan et al. [23] |
|  | H3K27AC | Link et al. [47] |
| fibroblast | ATACseq | Brumbaugh et al. [9] |
|  | H3K4ME1 | ENCODE |
|  | H3K4ME3 | ENCODE |
|  | H3K27AC | ENCODE |
| B | ATACseq | Mostafavi et al. [52] |
|  | H3K4ME1 | Mostafavi et al. [52] |
|  | H3K4ME3 | Daniiel et al. [19] |
|  | H3K27AC | Sabo et al. [67] |
| HeLa | ATACseq | Cho et al. [14] |
|  | H3K4ME1 | ENCODE |
|  | H3K4ME3 | ENCODE |
|  | H3K27AC | ENCODE |
| K562 | ATACseq | Antony et al., unpublished |
|  | H3K4ME1 | ENCODE |
|  | H3K4ME3 | ENCODE |
|  | H3K27AC | ENCODE |
| THP1 | ATACseq | Phanstiel et al. [59] |
|  | H3K4ME1 | Mohaghegh et al. [51] |
|  | H3K4ME3 | Nurminen et al. [55] |
|  | H3K27AC | Nurminen et al. [55] |

**Table 2:** STAT1, STAT2, and IRF9 ChIPseq Datasets. For each cell type, we downloaded ChIPseq datasets collected at both homeostasis and after IFN stimulation. For BMDM, fibroblast, HeLa, and THP1, we downloaded STAT1, STAT2, and IRF9 ChIPseq datasets. For B and K562 cells, only STAT1 and STAT2 ChIPseq datasets were available. Datasets varied in IFN subtype, hour of sampling post IFN stimulation, and IFN concentration as shown. See Supplementary Information for accessions and number of replicates.

| cell type | reference | details |
| --- | --- | --- |
| BMDM | Platanitis et al. [61] | IFNb, 1.5 hr, 250 IU/ml |
| fibroblast | Platanitis et al. [61] | IFNb, 1.5 hr, 250 IU/ml |
| B | Mostafavi et al. [52] | IFNa, 1.5 hr, 10,000 IU/ml |
| HeLa | Au-Yeung et al. [1] | IFNa, 2 hr, 1000 U/ml |
| K562 | ENCODE | IFNa, 6 hr, N/A |
| THP1 | Platanitis et al. [61] | IFNb, 1.5 hr, 250 IU/ml |

**Table 3:**
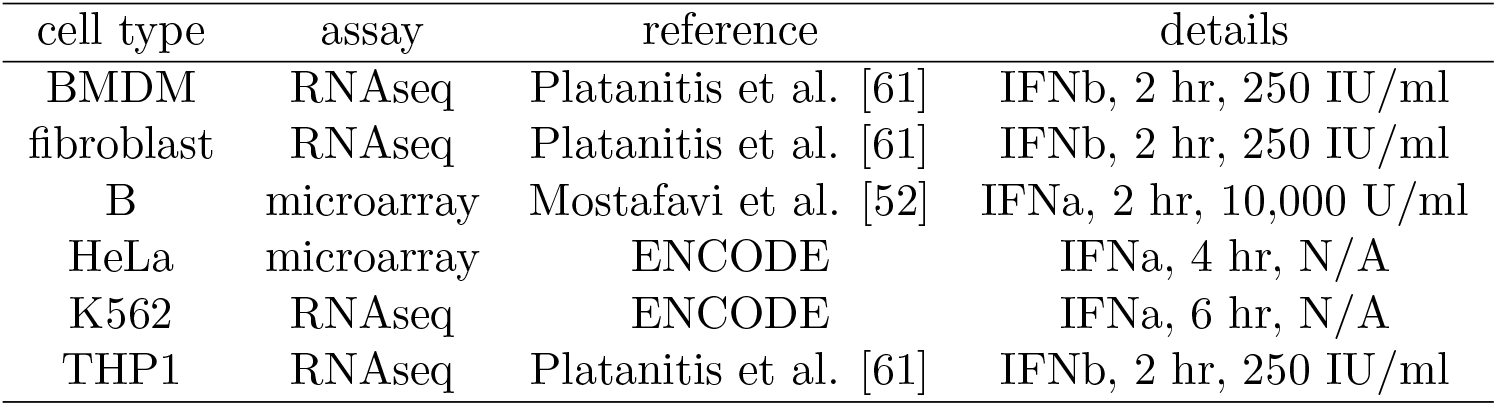
Transcription Datasets. For each cell type, we downloaded either RNAseq or mi-croarray datasets collected at both homeostasis and after IFN stimulation. Datasets varied in IFN subtype, hour of sampling post IFN stimulation, and IFN concentration as shown. See Supplementary Information for accessions and number of replicates.

### 2.1 Processing ATACseq and ChIPseq Datasets

For each cell type, we downloaded ATACseq and H3K4ME1, H3K4ME3, and H3K27AC ChIPseq datasets, all collected at homeostasis. For each cell type, we downloaded STAT1, STAT2, and IRF9 ChIPseq datasets at homeostasis and under IFN stimulation, except that for B and K562 cells only STAT1 and STAT2 ChIPseq datasets were available and for K562 cells no homeostatic STAT1 and STAT2 ChIPseq datasets were available; instead we downloaded K562, STAT1 and STAT2 datasets collected 30 minutes after IFN stimulation as an alternative for the homeostatic state.

For the NCBI datasets, we downloaded fastq files from the SRA dataset, used bowtie 1.2 [42] to align reads to the mouse mm10 genome and human hg38 genome, and then used MACS2, [87], to call peaks and form signals (SPMR). The number of replicates varied from 1 − 4 across the datasets and when multiple replicates were available we aligned each separately using bowtie, combined the bam files, and then used MACS2 to call peaks. We used the default settings in bowtie 1.2 and filtered the output bam file for mapped and, when appropriate, paired reads, and removed duplicate reads. We used the default settings in MACS for ChIPseq data, but for ATACseq added the options `--nomodel --shift -37 --extsize 73`. We called peaks at an FDR of 0.01. We used the MACS2 `--SPMR` option to create signal output. For ENCODE datasets, we downloaded ENCODE constructed peak calls and signal files, choosing the files constructed over all replicates.

### 2.2 Processing Transcription Datasets

For each cell type, we downloaded transcription datasets sampled at homeostasis and under IFN stimulation from GEO or ENCODE. Within each cell type, the homeostatic and IFN stimulated datasets were drawn from the same experiment, so that the protocols were identical except for the presence or absence of IFN stimulation. For K562 and B cells, we downloaded microarray datasets; for the other cell types we downloaded RNAseq datasets.

For each gene in the mm10 and hg38 genomes, we identified the transcript with highest expression level under IFN stimulation in the mouse BMDM dataset and in a human monocyte derived macrophage dataset (Andrade et al, unpublished, GSE125352). This gave us one transcript for every gene on which we performed differential expression analysis. We did not use the human macrophage dataset in our full analysis because we could not find STAT1, STAT2, and IRF9 ChIP datasets for that cell type, but we chose it to match the mouse BMDM dataset in cell type.

The microarray datasets were all Affymetrix arrays. We downloaded raw CEL files, processed the CEL files using the R oligo package [11] using the rma algorithm and quantile normalization, and then used the R limma package [63] to produce fold-change and FDR values for the differential expression of each transcript. For the RNAseq datasets, we downloaded raw fastq files, aligned them to the mm10 or hg38 transcriptome using Salmon 0.99 [58] with default setting except the addition of the flag --validateMappings. We normalized the Salmon, expression values using the R edgeR package’s [66] calcNormFactor function with the method parameter set to TMM, processed the output using the voom function in the limma package, and used the R limma package to calculate differential expression fold-change and FDR just as for microarray data.

We defined a gene to be an ISG if the gene’s expression level changed by more than 50% under IFN stimulation and if the FDR adjusted p-value for differential expression computed by the R limma package was less than 0.05 for the RNAseq datasets and 0.01 for the microarray datasets. The lower p-value cutoff for microarray datasets removed noise associated with low level transcripts, a known limitation of microarrays [79].

### 2.3 Associating Peaks with ISRE

We located every 10 base pair sequence across the mm10 and hg38 genomes with at most one nucleotide difference from the ISRE TTTCNNTTTC motif, considering both positive and negative strands. The collection of these 10 base pair sequences formed our ISREs. We then associated a STAT1, STAT2, or IRF9 peak, from a ChIP dataset under IFN stimulation, with an ISRE if the summit of the peak, as called by MACS2, was within 10 base pair of the 5’ or 3’ nucleotides of the ISRE. These associations gave us the different ISRE classes as described in the text, e.g. STAT2.IRF9 ISRE class.

For each ISRE, we considered the 1010 base pair window centered on the 10 base pair ISRE, and associated the ATACseq and ChIPseq SPMR values and the nucleotide sequence for those 1010 base pairs with the ISRE. We converted signals and nucleotide sequences to features (a collection of numeric values) as described in the text.

### 2.4 Classification

We built classifiers using L1 logistic regression implemented using the R glmnet package [28]. We always implemented a binary classification, involving two ISRE classes. We used the features formed from the ATACseq, ChIPseq, or sequence data to predict the class type of ISRE. Given a collection of ISRE loci to classify, we trained the classifier using 50% of the loci. To choose the L1 penalty parameter, we applied 10 fold cross validation to the training data and using maximization of AUROC (AUC) to select the parameter. We then tested our classifier on the held out data, using AUC and weighted precision at a fixed recall of either 25% or 50% as a measure of predication accuracy. Weighted precision was calculated by weighting loci so that the sum of weights of each ISRE class were equal. We used the R ROCR package [69] to compute AUC and precision values.

We restricted classification to pairs of ISRE classes with samples sizes sufficiently large so that an AUC greater than 0.55 and precision greater than 0.60 were statistically significant. To determine sample sizes that were sufficiently large, we permuted the ISRE class value while keeping the homeostatic signal fixed. We did this for each pair of ISRE classes we were classifying, computed AUC and precision for the permuted data, and then considered classifications for which the permuted AUC and precision were under 0.55 and 0.60, respectively, with 95% confidence.

### 2.5 GO Enrichment Analysis

We used the R package gprofiler2 [62] to perform GO enrichment analysis. Given a collection of genes, we called the gprofiler2 gost function, filtered for GO classes with less than 500 genes, and sorted the remaining classes in descending order of p-values.

## 3 Results

### 3.1 ISRE Signatures

From the GEO and ENCODE databases [5, 21], we downloaded STAT1, STAT2, and IRF9 ChIPseq datasets collected under IFN stimulation for mouse bone-marrow derived macrophages (BMDM), fibroblasts, and B cells, and in human HeLa, K562, and THP1 cells. For B and K562 cells, only STAT1 and STAT2 ChIPseq datasets were available. All datasets were collected between 1.5 − 6 hours post IFN stimulation, relatively early in the IFN response but during the time period shown to have the most ISGs upregulated [52]. Datasets did vary in the type I IFN, IFNa or IFNb, and the IFN concentration, see Table 2 for details.

For each ChIPseq dataset, we called peaks using MACS2, [87], restricting our attention to peaks within 100kb of gene transcription start sites (TSS) on the mouse (mm10) and human (hg38) genomes. BMDM had many more peaks than the other cell types, roughly 40, 000 versus 1500 to 6000, and mouse cell types had more peaks than human cell types, even when BMDM where excluded, see Table 4 for specific values. The large number of BMDM peaks may reflect the unique role of BMDM in innate response. The difference in peak counts between mouse and human cell types may be due to low sensitivity of ChIP antibody binding in human cells, particularly IRF9, as discussed in [61].

**Table 4:** Association of STAT1, STAT2, and IRF9 ChIPseq Peaks with ISRE. For each cell type, shown are (STAT1, STAT2, IRF9) the number of peaks called for the STAT1, STAT2 and IRF9 datasets, respectively, and (STAT1 (%), STAT2 (%), IRF9 (%)) the fraction of peaks that we associated with an ISRE. We associated a peak with an ISRE if the peak summit was within 10 base pair of the ISRE.

| cell type | STAT1 | STAT2 | IRF9 | STAT1(%) | STAT2(%) | IRF9(%) |
| --- | --- | --- | --- | --- | --- | --- |
| BMDM | 7577 | 17467 | 14738 | 0.50 | 0.48 | 0.56 |
| fibroblast | 59 | 1582 | 2251 | 0.17 | 0.67 | 0.70 |
| B | 646 | 5647 |  | 0.19 | 0.33 |  |
| HeLa | 261 | 794 | 622 | 0.35 | 0.59 | 0.68 |
| K562 | 676 | 1408 |  | 0.53 | 0.40 |  |
| THP1 | 825 | 2888 | 164 | 0.08 | 0.11 | 0.18 |

#### ChIPseq Peaks and ISRE

In order to associate ChIP peaks with ISRE, we defined an ISRE as any 10 base pair sequence that matched the canonical ISRE motif TTTCNNTTTC or differed from it by a single nucleotide (where NN allowed for any nucleotide) and then located all such sequences within 100kb of a gene TSS on either the positive or negative strands. We associated an ISRE with a ChIP peak if the ISRE was within 10 base pairs of the peak summit.

Under our definition for ISREs, there are 2.5 and 3.1 million ISRE within 100kb of TSS on the mouse and human genome, respectively, but only several thousand were associated with peaks. Conversely, while few ISRE associated with peaks, a large percentage of peaks associated with ISRE as shown in Table 4. Except for THP1 cells, greater than 30% and 56% of STAT2 and IRF9 peaks, respectively, associated with an ISRE. Between 34% − 50% of STAT1 peaks associated with ISRE for BMDM, HeLa, and K562, but less than 20% for fibroblast and B cells. Since sequences that vary by more than one nucleotide from TTTCNNTTTC may serve functionally as ISREs, a higher fraction of peaks may have associated with ISRE than our estimates.

For a given cell type, we refer to an ISRE associated with a peak as a *bound ISRE* and we classify bound ISRE by the combination of associated STAT1, STAT2, and IRF9 peaks: the STAT2 ISRE class are those ISRE associated with just STAT2 peaks, the STAT2.IRF9 ISRE class are those ISRE associated with both a STAT2 and an IRF9 peak, but not a STAT1 peak, and so on. Across 3 of the 4 cell types for which we had STAT1, STAT2, and IRF9 ChIP data - namely BMDM, fibroblasts, and HeLa cells, with THP1 left out - 99% of bound ISRE were in the STAT2 (bound by just STAT2), STAT2.IRF9 (bound by STAT2 and IRF9), STAT1.STAT2.IRF9 or IRF9 ISRE classes, and associations involving STAT1 in the absence of STAT2 were rare. Further, in all three cell types, greater than 60% of bound ISRE were in the STAT2.IRF9 or STAT1.STAT2.IRF9 ISRE classes. In contrast, in THP1, ISRE in the STAT1 (just STAT1) or STAT2 (just STAT2) classes accounted for 11% and 77%, respectively, of bound ISRE, possibly reflecting sensitivity issues in ChIP antibody binding. In K562 and splenic B cells, the two cell types for which we did not have IRF9 ChIP data, 99% of bound ISRE were in the STAT2 or STAT1.STAT2 ISRE classes, in line with the large percentage of bound ISRE in the STAT2, STAT2.IRF9 and STAT1.STAT2.IRF9 classes in BMDM, fibroblast, and HeLa cells. See Table 5.

**Table 5:** The Number of Bound ISRE and their Distribution across ISRE Classes. We associated STAT1, STAT2, and IRF9 peaks with ISRE as described in the text. We referred to ISRE associated with at least one peak type as bound ISRE. For each cell type, shown are (n) the number of bound ISRE and the percentage of bound ISRE in each ISRE class. For B and K562 cells, we did not have IRF9 ChiPseq datasets, limiting us to STAT1, STAT2, and STAT1.STAT2 ISRE classes. (S1.S2=STAT1.STAT2, S2.I9=STAT2.IRF9, and so on).

| cell type | n | STAT1 | STAT2 | IRF9 | S1.S2 | S1.I9 | S2.I9 | S1.S2.I9 |
| --- | --- | --- | --- | --- | --- | --- | --- | --- |
| BMDM | 11818 | 0.00 | 0.09 | 0.08 | 0.01 | 0.01 | 0.40 | 0.41 |
| fibroblast | 3287 | 0.00 | 0.04 | 0.36 | 0.00 | 0.00 | 0.59 | 0.00 |
| HeLa | 890 | 0.01 | 0.18 | 0.16 | 0.00 | 0.00 | 0.51 | 0.14 |
| THP1 | 509 | 0.11 | 0.77 | 0.03 | 0.01 | 0.00 | 0.04 | 0.04 |
| B | 2760 | 0.01 | 0.94 |  | 0.05 |  |  |  |
| K562 | 874 | 0.01 | 0.35 |  | 0.64 |  |  |  |

Overall, our analysis of peaks and ISRE is consistent with existing results [6, 61]: most binding of ISGF3 components was associated with ISREs, STAT1 binding to ISRE independent of STAT2 was rare, and most ISRE binding involved STAT2:IRF9 dimers or ISGF3.

#### ISRE Binding Across Genes

We associated each bound ISRE to the gene with the closest TSS. Since promoter and enhancer regulatory elements differ in chromatin state and functional properties [22], we split the region within 100kb base pairs of a gene’s TSS into an enhancer region that included positions 3 − 100kb from the TSS and a promoter region that included positions within 500 base pairs of the TSS. We do not claim that ISRE in the enhancer and promoter regions act functionally as enhancers or promoters, respectively; instead, we use the terminology putatively and to emphasize the different characteristics of ISRE in the two regions, as we describe below. We purposely left a gap, positions 500 − 3000 from the TSS, between the promoter and enhancer regions for greater separation of chromatin state and functionality.

We were interested in the number of bound ISRE and their positional distribution within genes. While we could identify bound ISRE with high specificity (given that a bound ISRE reflected the intersection of an ISRE and a ChIPseq peak), we likely missed bound ISRE that were not within one nucleotide of the TTTCNNTTTC motif, as noted above. With this in mind, we used the absence of STAT1, STAT2, and IRF9 peaks in a region, rather than the absence of bound ISRE, as evidence that no bound ISRE existed in the region.

Of genes with at least one bound ISRE, which we call *bound genes*, 82% in mouse cell types and 95% in human cell types had between 1 − 6 bound ISRE. A small number of bound genes had a large number of bound ISRE, reflecting repeating, adjacent ISRE. For example, in BMDM, the gene RHOH had 16 ISRE contained in a region of 185 base pairs (chr5:65834014-65834198 in mm10) of which 13 were in the STAT1.STAT2.IRF9 bound ISRE class. Figure 1 shows the number of bound ISRE in BMDM at different distances relative to the TSS. Other cell types showed a similar pattern. Bound ISRE were at their higher concentration near the TSS, in the promoter region, and fell in concentrations with distance from the TSS. However, most bound ISRE, 82% and 70% in mouse and human cell types respectively, fell in the enhancer region.

**Figure 1:**
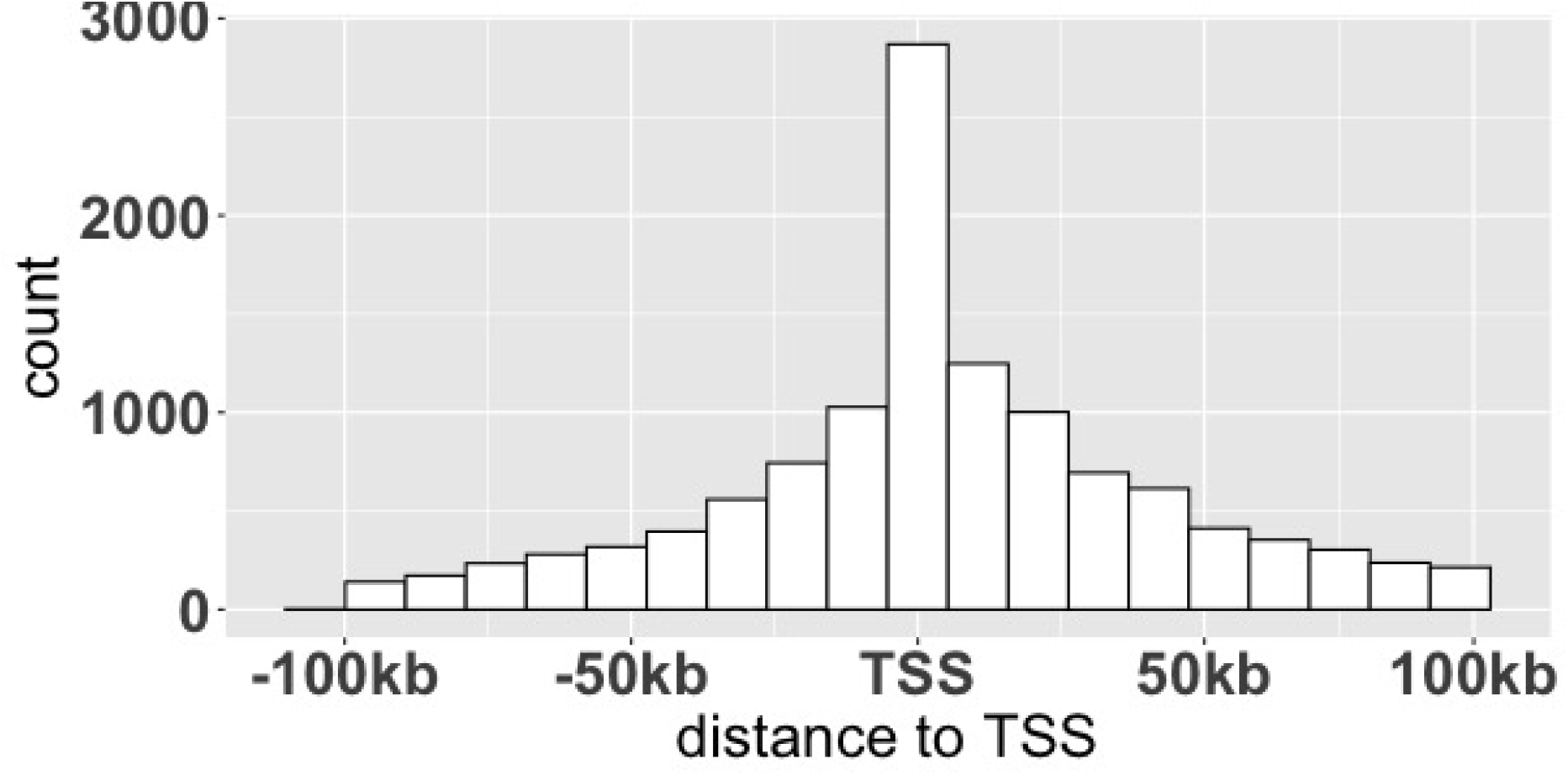
The Distribution of ISREs Relative to TSS. Shown are the counts of bound ISRE in BMDM according to the distance of the ISRE to the gene transcription start site (TSS). Bound ISREs were most dense near the TSS, in promoter regions, but 70% − 80% of bound ISRE were distal to the TSS, in enhancer regions.

To understand the positional distribution of bound ISRE within genes, we considered three configurations of bound genes: a bound ISRE in the promoter region, in the enhancer region, and in both the promoter and enhancer region. Across all cell types, between 12 − 25% of bound genes had a bound ISRE in the promoter while greater than 60% of bound genes had a bound ISRE in the enhancer region. Bound genes with bound ISRE in both the enhancer and promoter were rare, less than 5% across all genes

Since we may have missed a substantial number of bound ISRE, the percentages of these three configurations may be biased. To address this potential bias, we considered two additional positional configurations: a bound ISRE just in the promoter and a bound ISRE just in the enhancer. For these two configurations, we used the absence of a STAT1, STAT2, or IRF9 peak to infer the absence of a bound ISRE as discussed above. Across cell types, greater than 43% of bound genes had a bound ISRE just in the enhancer. And, across all cell types except BMDM, greater than 11% of bound genes had an ISRE just in the promoter, with BMDM having 3%. See the upper panel of Figure 2.

**Figure 2:**
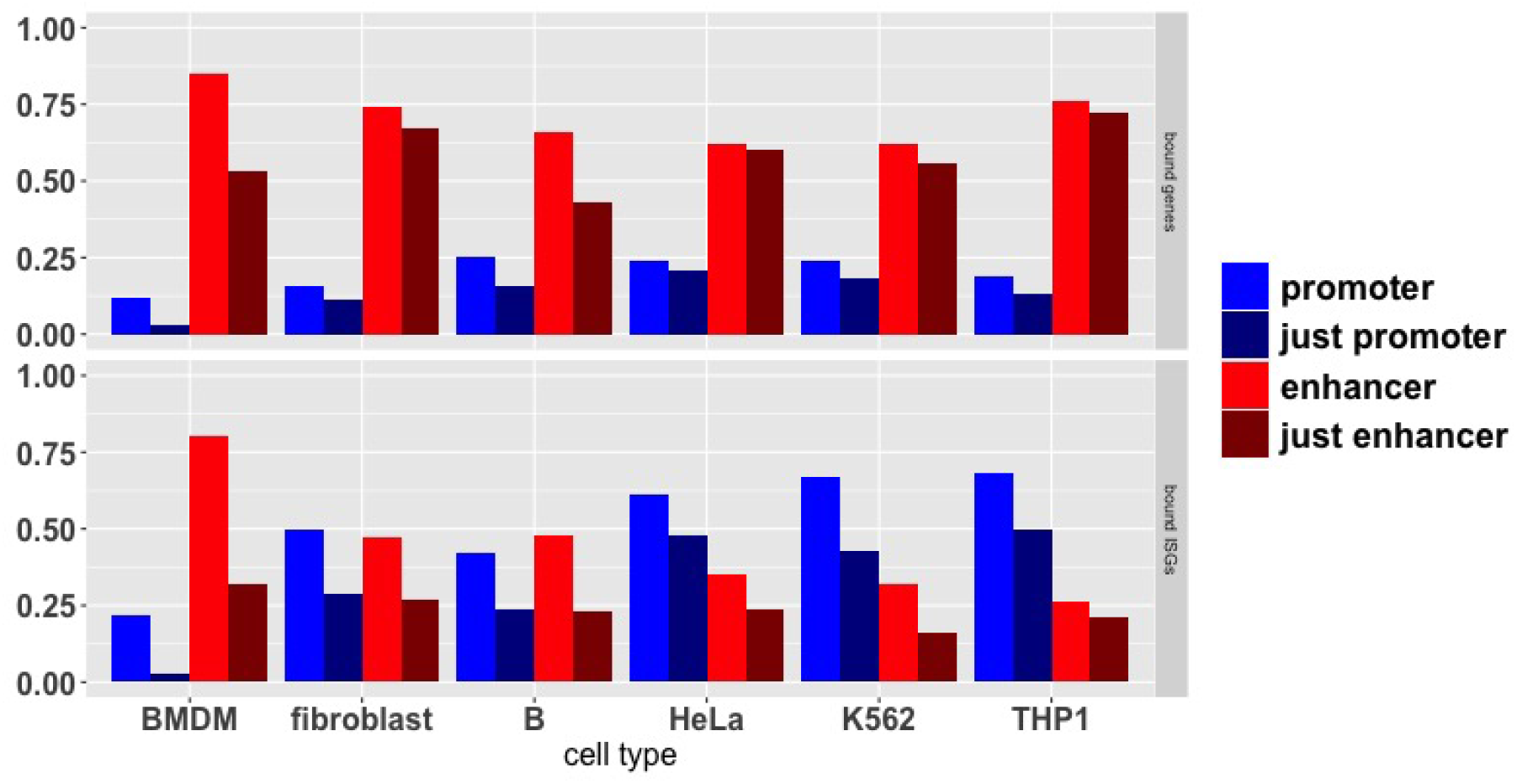
The Distribution of Bound ISREs Across Promoter and Enhancer Regions. We defined a bound gene as a gene with a bound ISRE within 100kb of the TSS. (Upper panel) For each cell type, bars show (left to right) the fraction of bound genes with a bound ISRE in the promoter region, with bound ISRE just in the promoter region, with a bound ISRE in the enhancer region, and with a bound ISRE just in the enhancer region. (Lower panel) Same as the upper panel except that we consider bound ISGs rather than bound genes.

These results support a model in which a substantial percentage of ISRE binding is restricted to the enhancer regions of a gene. However, our ability to rule out the presence of a bound ISRE through the absence of peaks depends on the aggressiveness of our peak calling, which we set using an FDR of 0.01 in MACS2. More aggressive peak calling would result in more peaks and therefore reduce the percentage of bound genes that we place in the just enhancer category. But peak calling is dependent on the ChIP signal levels, and if we are missing peaks they are necessarily peaks with lower signal. In this case our results would switch from a dichotomy, i.e. a substantial portion of bound genes have bound ISRE only in the enhancer region, to a continuum, i.e. a substantial portion of bound genes have strongly bound ISRE only in the enhancer region. Regardless of whether peaks are present or absent in promoter regions, our results do show that most bound genes have bound ISRE in their enhancer regions.

#### ISRE Binding Across Cell Types

We next considered the degree to which bound ISRE were shared across cell types. Roughly 90% of bound ISRE in fibroblasts and B cells were bound in BMDM, while about 20% of BMDM bound ISRE were bound in the other two cell types. Fibroblast and splenic B cells shared about 25% of their bound ISRE, giving an overall picture of broad ISRE binding in BMDM with fibroblasts and splenic B binding reflecting different subsets of the ISRE bound in BMDM. Splitting bound ISRE by enhancer and promoter region, we found that 20% of bound ISRE in the enhancer region were shared across all three cell types while 60% of bound ISRE in the promoter region were shared. Overall, 578 and 312 bound ISRE located in the enhancer and promoter regions, respectively, were shared across all three cell types. See Table 6 for more details

**Table 6:** Overlap of Bound ISRE Across Cell Types. Shown are the fraction of bound ISRE shared in promoter regions (top tables) and enhancer regions (bottom tables) in mouse cell types (left tables) and human cell types (right table). The diagonal of each table gives the fraction of bound ISRE in the cell type that are also bound in both of the other two cell types. The off diagonals give the fraction of bound ISRE in the row cell type that are also bound in the column cell type. Bound ISRE in promoter regions are shared at higher frequencies than bound ISRE in enhancer regions. (FB=fibroblast)

| promoter | BMDM | FB | B | promoter | HeLa | K562 | THP1 |
| --- | --- | --- | --- | --- | --- | --- | --- |
| BMDM | 0.41 | 0.5 | 0.6 | HeLa | 0.54 | 0.9 | 0.55 |
| FB | 0.97 | 0.79 | 0.8 | K562 | 0.84 | 0.5 | 0.53 |
| B | 0.88 | 0.61 | 0.6 | THP1 | 0.92 | 0.97 | 0.92 |
| enhancer |  |  |  | enhancer |  |  |  |
| BMDM | 0.06 | 0.22 | 0.17 | HeLa | 0.09 | 0.43 | 0.11 |
| FB | 0.89 | 0.23 | 0.24 | K562 | 0.46 | 0.1 | 0.13 |
| B | 0.88 | 0.32 | 0.31 | THP1 | 0.17 | 0.18 | 0.14 |

For human cell types, there was no cell type with a broader response than the other two. Instead, HeLa and K562 shared roughly 60% of their bound ISRE while THP1 shared 20% of bound ISRE with each of the other cell types. THP1 had about 500 bound ISRE while the other two cell types both had roughly 900, with the difference possibly due to the already mentioned technical issues with the ChIP datasets. As in mouse cell types, bound ISRE in the enhancer region were shared across all cell types at a lower percentage than bound ISRE in the promoter region, 11% and 65% respectively. Overall, 51 and 108 bound ISRE in the enhancer and promoter regions, respectively, were shared across all three cell types.

We performed a GO ontology enrichment analysis ([16, 50]) on genes with bound ISRE that were shared across cell types, splitting the analysis into genes with shared bound ISRE in promoter and enhancer regions, respectively; see Methods for details. For promoter regions, 169 genes had bound ISRE shared across all three mouse cell types. The three GO classes with the highest p-value were *response to virus, defense response to virus, and response to interferon-beta* and 33 of the 169 genes, 20%, were in the *defense response to virus* class. In contrast, 297 genes shared bound ISRE in the enhancer region and the three GO classes with the highest p-value were *regulation of immune effector process, regulation of hemopoiesis, and regulation of chromatin binding*. 15 of the 297 genes were in the *defense response to virus* class, 5%, significantly less than what we found for promoter genes (p-value 5E-13).

GO enrichment analysis for the human cell types gave similar results. Sixty genes shared bound ISRE in the promoter region with the top GO ontology classes of *defense response to virus, response to virus, and response to type I interferon*. 27 of 60 genes, 45%, were classified in the *defense response to virus* GO class. For the enhancer region, 29 genes shared bound ISRE in the enhancer region across cell types. The top three enriched GO classes were *cellular response to type I interferon, type I interferon signaling pathway, and response to type I interferon*. 5 of the 29 genes, 17%, were in the *defense response to virus*, significantly less than for the promoter region (p-value 0.001).

Overall, genes with shared bound ISRE in the enhancer region were labeled as antiviral at significantly lower percentages than genes in the promoter region. This difference may reflect functional differences, but the results may reflect gene annotations derived from IFN studies which have tended to concentrate on genes with ISRE in the promoter region [20]. Overlap between bound ISRE in the enhancer region, 6% *−*31%, was lower than in promoter regions, 41% − 92%, in line with existing models of cell-type specificity mainly associated with enhancer variation [33, 34, 17]. The high percentage of overlapping bound ISRE in promoter regions are especially significant given the disparate cell types considered, e.g. fibroblasts vs. B cells and HeLa cells vs. K562 cells. Overall, these results suggest a model in which binding of ISRE in promoter regions is largely conserved across cell types and associates with effectors of viral defense, while binding of ISRE in enhancer regions controls cell-specific components of the IFN response.

### 3.2 ISRE Binding and Differential Expression

To investigate the correlation between bound ISRE and ISGs, we downloaded transcription datasets collected under homeostasis and under IFN simulation: RNAseq datasets for BMDM, fibroblast, HeLa, and THP1 cells and microarry datasets for B and K562 cells. As was the case for the ChIPseq datasets, the datasets varied in IFN subtype (IFNa or IFNb), the concentration of IFN, and the time of sampling, which varied between 2 − 6 hours. For all the cell types, the transcription datasets and the STAT1, STAT2, and IRF9 ChIPseq datasets collected under IFN stimulation were exposed to the same IFN subtype and concentration, although the time of sampling varied slightly, see Tables 2 and 3 for specific values

We defined a gene to be an ISG, meaning that it is differentially expressed under IFN stimulation, if the gene’s expression level changed by more than 50% under IFN stimulation and if the FDR adjusted p-value for differential expression was less than 0.05 for the RNAseq datasets and 0.01 for the microarray datasets, see Methods for details. BMDM had roughly 1700 ISGs, consistent with the larger number of bound ISRE and the central role of BMDM in the innate response. The other cell types had roughly 400 ISGs, except for B cells, which had roughly 700, possibly reflecting a broader interferon response in immune cells. In mouse cell types (BMDM, fibroblast and B cells), between 40% − 62% of ISGs were bound, i.e. had a bound ISRE within 100kb of the TSS. In human cell types (HeLa, K562 and THP1), the fraction was much lower, 9% − 15%, see Table 7. In THP1, the already noted sensitivity issue may explain the small percentage, and lack of sensitivity may extend to the other cell types as well. Regardless of the exact percentages, across all cell types a substantial percentage of ISGs were bound. That some ISGs were unbound may reflect bound ISRE missed by our analysis, particularly in the case of the human cell types, or interferon signaling pathways independent of ISGF3 components.

**Table 7:** Bound ISGs. We defined a bound ISG as an ISG with a bound ISRE within 100kb of the TSS. Shown are the (ISG) number of ISGs, (bound ISG) the number of bound ISGs, (bound ISG (%)) and the fraction of ISGs that are bound.

| cell type | ISG | bound ISG | bound ISG (%) |
| --- | --- | --- | --- |
| BMDM | 1729 | 1083 | 0.63 |
| fibroblast | 493 | 214 | 0.43 |
| B | 378 | 205 | 0.54 |
| HeLa | 668 | 66 | 0.10 |
| K562 | 415 | 63 | 0.15 |
| THP1 | 444 | 68 | 0.15 |

We next investigated the position distribution of bound ISRE within bound ISGs. The lower panel of Figure 2 shows the fraction of bound ISGs with bound ISRE in promoter regions, enhancer regions, and just promoter and just enhancer regions, and is analogous to the top panel of Figure 2, which gives the same fractions but as a total of bound genes rather than bound ISGs. Comparison of the lower and upper panels of Figure 2 shows that bound ISGs are enriched for bound ISRE in their promoters relative to bound genes. However, a substantial percentage of bound ISGs had bound ISRE in their enhancer region and just in their enhancer region, although the aggressiveness of our peak calling might bias the percentage of ISGs with bound ISRE just in the enhancer regions, as previously mentioned. Depending on the cell type, 26% − 80% and 16% − 32% of bound ISGs had a bound ISRE in the enhancer region and just in the enhancer region, respectively.

To test for a correlation between ISRE binding and ISGs, we compared the fraction of genes that were ISGs under different positional configurations of bound ISRE. Of genes with a bound

ISRE in the promoter region or in the enhancer region, 22% − 68% and 26% − 80% were ISGs, respectively. To deconvolve the contribution of bound ISRE in promoters and enhancer regions, we compared genes with no ChIP peaks to genes with a bound ISRE just in the enhancer, just in the promoter, and in the enhancer and promoter, respectively; see Figure 3. Of genes with no peak, between 1% − 3% were ISGs, depending on the cell type. For all cell types except THP1, the percentage of genes that were ISGs rose as we moved from no peaks to a bound ISRE just in the enhancer to a bound ISRE just in the promoter to a bound ISRE in the enhancer and promoter. When a bound ISRE was just in the enhancer, 2% − 12% of genes were ISGs, modestly above, and statistically significantly greater in all cell types except K562, than the 1% − 3% seen in genes with no peak. When a bound ISRE was just in the promoter, 25% − 69% of genes were ISGs, significantly above the percentages for bound ISRE just in the enhancer across all the cell types. Relatively few genes had bound ISRE in the promoter and enhancer, limiting our statistical power. With that in mind, when a bound ISRE was in both promoter and enhancer, 33% − 100% of genes were ISGs, significantly greater than the percentage for just promoters in BMDM and fibroblast, but not the other cell types, possibly due to the small samples sizes.

**Figure 3:**
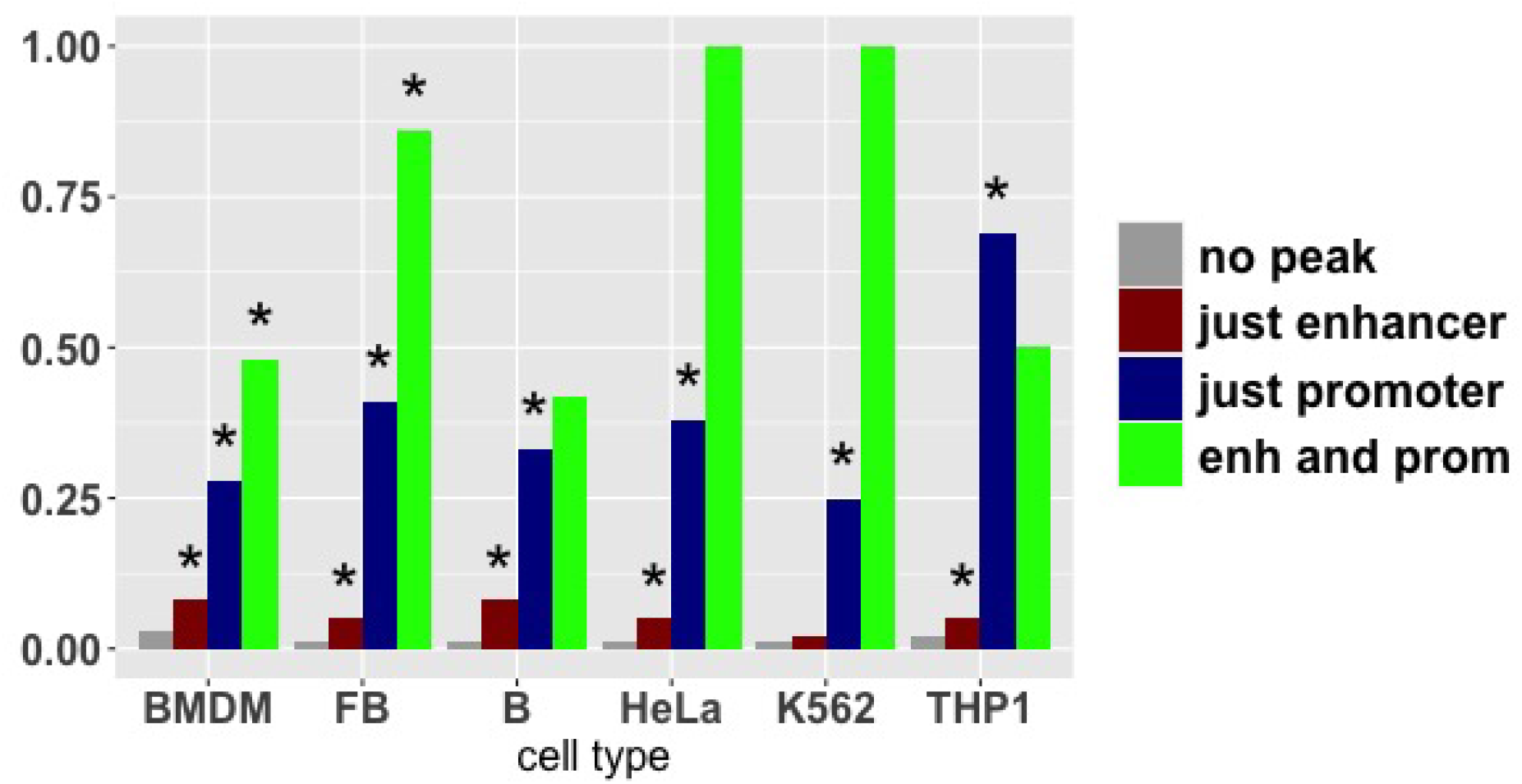
Associations Between Bound ISRE and ISGs. For each cell type, bars show (left to right) the fraction of genes that are ISGs over genes with (no peak) no STAT1, STAT2, and IRF9 ChIPseq peaks, (just enhancer) a bound ISRE just in the enhancer region, (just promoter) a bound ISRE just in the promoter region, and (enh and prom) bound ISRE in both the enhancer and promoter regions. Asterisks identify a statistically significant increase, at greater than 95% confidence, relative to the previous (bar to the left) frequency and configuration, see text for details. (FB=fibroblast)

We also considered the association between ISRE binding and the fold-change (FC) in expression from homeostasis to IFN stimulation. In ISGs with no peak, a bound ISRE just in the enhancer, a bound ISRE just in the promoter, and bound ISRE in both promoter and enhancer, the average FC across cell types was 1.4, 1.6, 2.6, and 3.0 log2, respectively.

We next considered the association of the STAT2, STAT2.IRF9, and STAT1.STAT2.IRF9 ISRE classes with ISGs. To compare expression across classes, we considered genes with a single bound ISRE either in the enhancer or in the promoter and no peak elsewhere. (Considering a single bound ISRE controlled for possible correlations between the ISRE class and the number of bound ISRE.) There was not sufficient sample size to consider genes with a single bound ISRE in both the promoter and enhancer regions. We necessarily restricted to cell types for which we had an IRF9 ChIP dataset, and we left out THP1 due to the sensitivity issues, leaving BMDM, HeLa and fibroblasts.

When the bound ISRE was in the promoter region, see the top panel of Figure 4, the percentage of genes that were ISGs rose sequentially as we considered the STAT2, STAT2.IRF9, and STAT1.STAT2.IRF9 ISRE classes, respectively. Under STAT2 binding, 5% − 12% of genes were ISGs, under STAT2:IRF9 binding, 18% − 46%, and under STAT1:STAT2:IRF9 binding, 58% − 60%. STAT2 binding percentages were not significantly greater than the 1% − 3% we noted in genes with no peak, supporting non-functional or weakly functional binding. STAT2.IRF9 percentages were significantly greater than those in genes with no peak in BMDM and fibroblast but not HeLa, and STAT1.STAT2.IRF9 percentages were significantly greater than STAT2.IRF9 percentages, suggesting a functional hierarchy.

**Figure 4:**
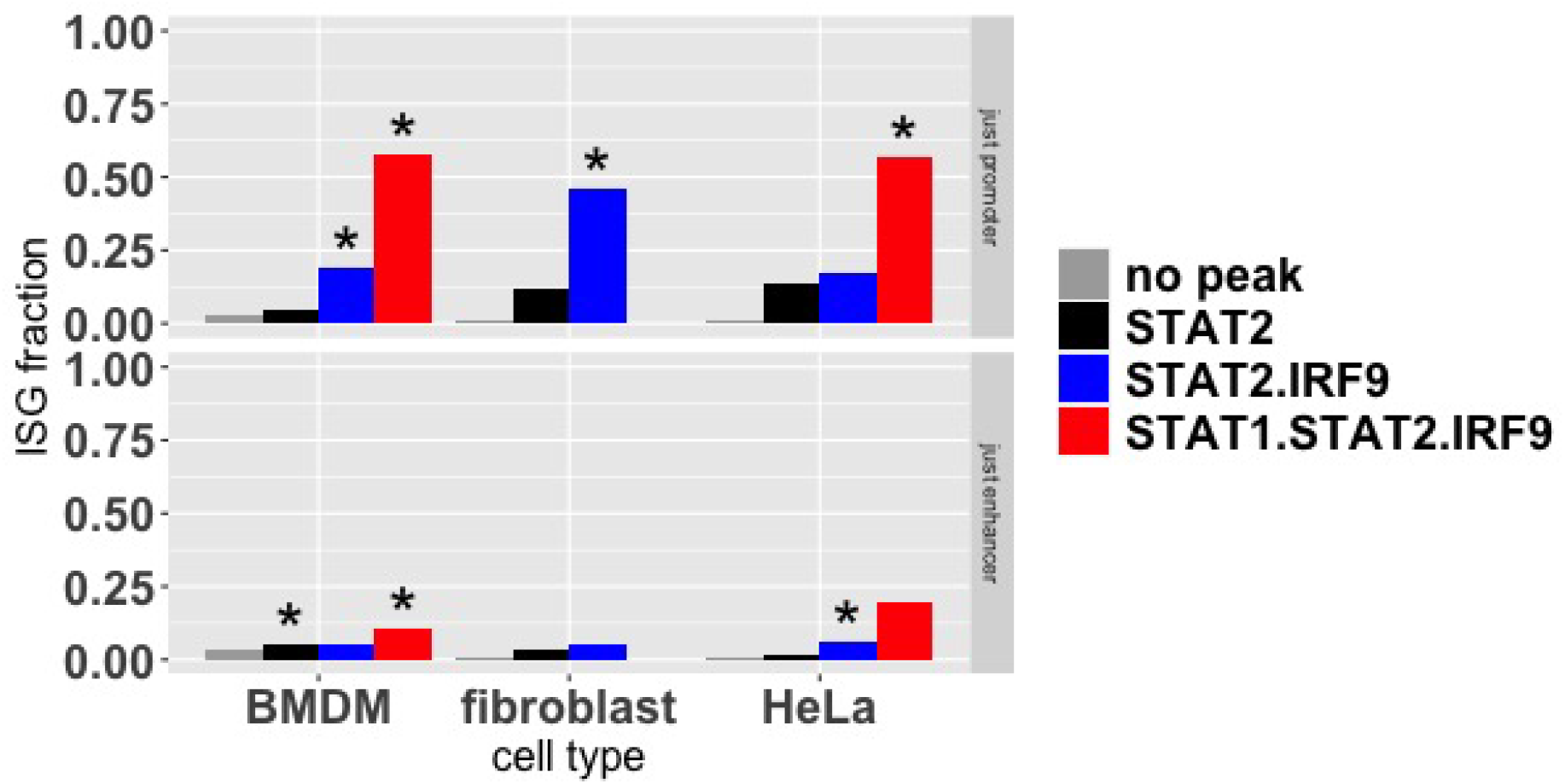
Associations Between Bound ISRE Class and ISGs. For each cell type, bars show (left to right) the fraction of genes that are ISGs over genes with no peak, a single bound ISRE of class STAT2, STAT2.IRF9, or a single STAT1.STAT2.IRF9 in just the promoter (top panel) or just the enhancer (bottom panel) regions. The percentage of genes that are ISGs increases according to ISRE binding by STAT2, STAT2 and IRF9, and STAT1, STAT2, and IRF9, suggesting a functional hierarchy. Asterisks identify a statistically significant increase, at greater than 95% confidence, relative to the previous (bar to the left) frequency and ISRE class.

When the bound ISRE was in the enhancer region, see the bottom panel of Figure 4, percentages also rose sequentially as we considered STAT2, STAT2.IRF9, and STAT1.STAT2.IRF9 ISRE classes, respectively, but overall the percentages were lower than when the binding was in the promoter region. For binding in the enhancer region, percentages associated with STAT2 and STAT2.IRF9 binding were not significantly greater than the percentage in genes with no peak. But 10% − 11% of genes bound by STAT1.STAT2.IRF9 were ISGs, which was significantly greater than genes with no peak.

ChIPseq binding levels associate with functionality [26, 45], so we considered the summit heights of STAT2 ChIP peaks associated with the different ISRE classes. As shown in Figure 5, STAT2 ISRE had significantly lower summit heights than the other classes, and STAT2.IRF9 had significantly lower summit heights than STAT1.STAT2.IRF9.

**Figure 5:**
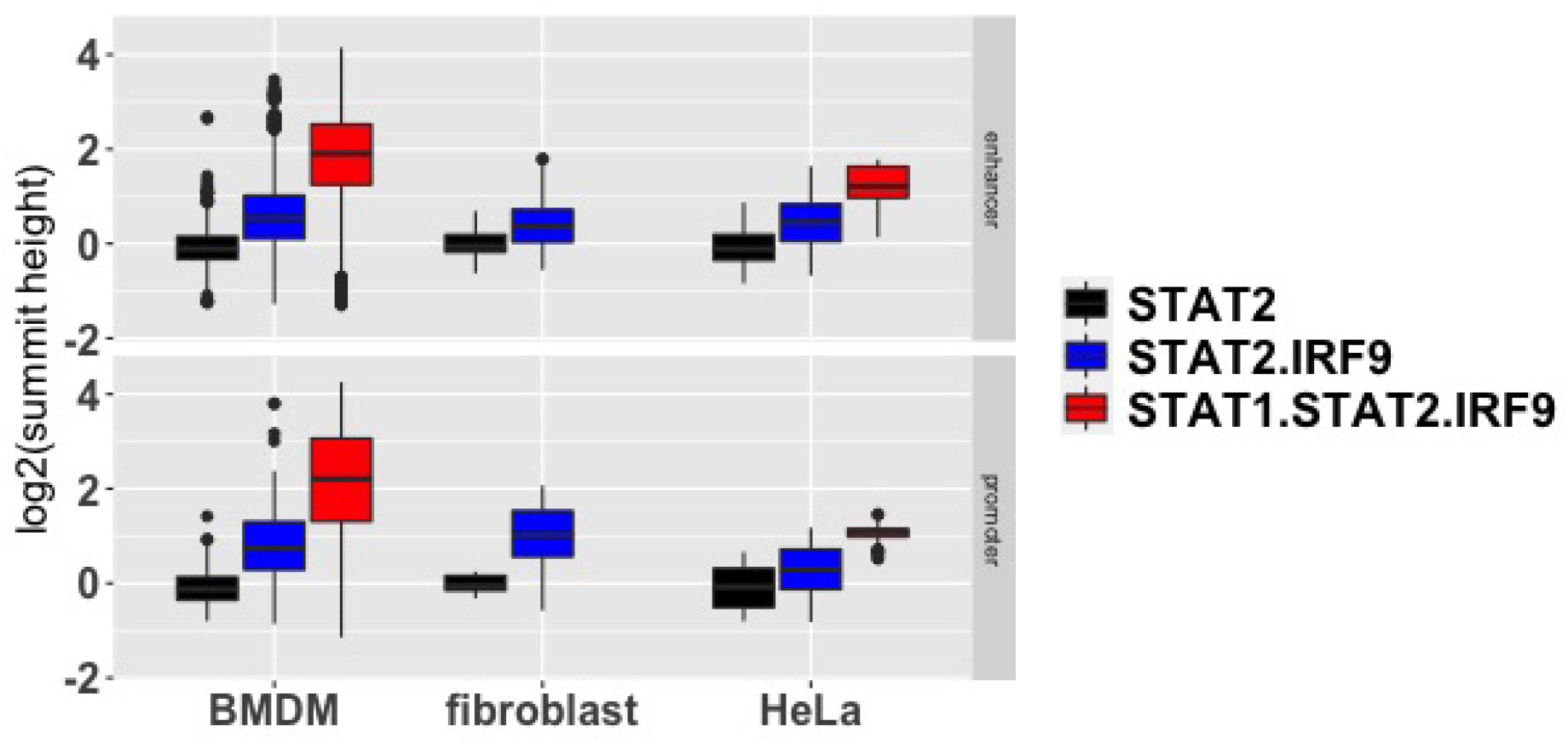
ChIPseq Binding Levels of STAT2 across ISRE Classes. Shown is the distribution of the summit heights for peaks in STAT2 ChIPseq datasets at ISRE of different classes. The STAT1.STAT2.IRF9 ISRE class summit heights were greatest, suggesting the strongest binding to ISRE and supporting a functional hierarchy across the ISRE binding classes.

Overall, our analysis shows a strong association between ISRE binding in the promoter region and differential expression under IFN stimulation. We also noted a significant, though weaker, association between ISRE binding in the enhancer regions and differential expression. However, the association is weak or absent when we restrict to ISRE bound by STAT2, and weaker in ISRE bound by STAT2.IRF9 than ISRE bound by STAT1.STAT2.IRF9. Our STAT2 ISRE class may represents misclassifications of STAT2.IRF9 or STAT1.STAT2.IRF9 ISRE. The inability of STAT2 to stably bind DNA in the absence of IRF9 supports this viewpoint [8] In contrast, multiple reports support the existence of STAT2.IRF9 dimers that are functionally important and independent of STAT1 expression [25, 6, 61].

### 3.3 Predicting ISRE Binding from Homeostatic, Epigenetic State

We next asked whether a cell’s homeostatic epigenetic state and DNA sequences associated with ISRE loci could be used to accurately predict ISRE binding under IFN stimulation. Associated with each ISRE, we formed a 1010 base pair locus composed of the 10 base pair ISRE motif and the 500 base pairs upstream and downstream of the motif. To describe the homeostatic, accessibility and histone state, we collected ATACseq and H3K4ME1, H3K4ME3, and H3K27AC ChIPseq datasets for each of our cell types. All datasets were downloaded from GEO or ENCODE and were collected at homeostasis, see Table 1 for details. For each ATACseq and ChIPseq dataset, we used MACS2 to build a signal (SPMR) over the 1010 bound locus, see Materials and Methods for details. To quantify the presence of co-transcription factor binding sites, we collected TF motifs from the JASPAR database [27] and matched them to the the ISRE loci sequences. To account for sequence specificities, we used the nucleotide sequence of the ISRE loci to describe the ISRE motif, CpG content, and DNA shape. Finally, we also downloaded STAT1, STAT2, and IRF9 ChIPseq datasets collected at homeostasis and formed signal for each ISRE locus from these datasets, as we did with the ATACseq and histone datasets, see Table 2. Overall, we characterized the homeostatic state through chromatin accessibility (ATACseq), histone markings (H3K4ME1, H3K4ME3, H3K27AC ChIPseq datasets), co-transcription factor motifs (JASPAR motifs), sequence specificities (ISRE locus DNA sequence), and STAT1, STAT2, and IRF9 binding (ChIPseq).

To quantify the predictive capacity of the homeostatic state, we built binary classifiers to distinguish pairs of ISRE classes based on the homeostatic data. We considered classes of bound ISRE and unbound ISRE. Importantly, these classes were defined using datasets collected under IFN stimulation. The bound ISRE classes included the classes we described above, reflecting the associations of STAT1, STAT2, and IRF9 with an ISRE - e.g. the STAT2 ISRE class, the STAT2.IRF9 ISRE class etc - but we also considered ISRE that were bound in the given cell type, at least by STAT2 but possibly also by STAT1 and IRF9, but were unbound in the other cell types, referring to these ISRE as the self ISRE class. We also defined classes of unbound ISRE: ISRE that were not bound by STAT1, STAT2, or IRF9 in any cell type, which we called the non-peak class, loci that were not associated with an ISRE, which we called the background class, and ISRE that were not bound by STAT1, STAT2, or IRF9 in the given cell type, but were bound in some other cell type, which we called the complement ISRE class. The unbound classes represent a hierarchy: loci in the background class don’t have an ISRE, loci in the non-peak class have ISREs that are not bound in any cell type, loci in the complement class are not bound in the given cell type but are bound in other cell types.

We built binary classifiers to distinguish a bound ISRE class from an unbound ISRE class in each cell type. For example, we built a classifier to distinguish the STAT1.STAT2.IRF9 ISRE class from the non-peak class in BMDM, meaning that we collected all ISRE loci in BMDM bound by STAT1.STAT2.IRF9 and all ISRE loci unbound in any cell type and built a classifier that distinguished the two ISRE classes based on the homeostatic state of ISRE loci. Given our results suggesting that the STAT2 ISRE class represents non-functional binding, we also built classifiers to distinguish the bound ISRE classes, other than the STAT2 ISRE class, from the STAT2 ISRE class. As described more fully below and in the Methods, we converted the homeostatic signals, JASPAR motifs, and nucleotide sequences of ISRE loci into features and used L1 logistic regression to build the binary classifiers. In all cases, we trained each classifier using half the relevant ISRE loci and used the other half to test prediction accuracy. We used AUROC (AUC) and precision as measures of prediction accuracy. In the main text, we present AUC values, precision values are provided in the Supplementary Information. We only built classifiers for pairs of classes with sufficient sample size (i.e. sufficient number of ISRE in each class) so that an AUC greater than 0.55 and a precision of greater than 0.60 was statistically significant at 0.05 level, see Materials and Methods for details.

We constructed the non-peak ISRE class through a one to one mapping to bound ISRE. As we mentioned above, there were 2 − 3 million non-peak ISRE (unbound ISRE motifs), but only a few thousand bound ISRE (12, 438 on the mouse genome and 1, 548 on the human genome, to be exact). For every bound ISRE, we randomly chose a non-peak ISRE with matching distance to a gene TSS, where the genes we considered where the collection of genes with bound ISRE. We built the background loci class in the same way, except that we randomly chose a locus rather than an ISRE motif. Importantly, by choosing the non-peak and background class in this limited way, we assessed our ability to distinguish a bound ISRE from a typical unbound ISRE, rather than our ability to distinguish a bound ISRE from all unbound ISRE.

#### Chromatin Accessibility

Figure 6A shows the mean homeostatic, ATACseq signal for ISRE, split by ISRE class, cell type and promoter or enhancer region. To normalize the signals, for each cell type and gene region combination, we scaled the signal so that the background class had a mean signal of 0.

**Figure 6:**
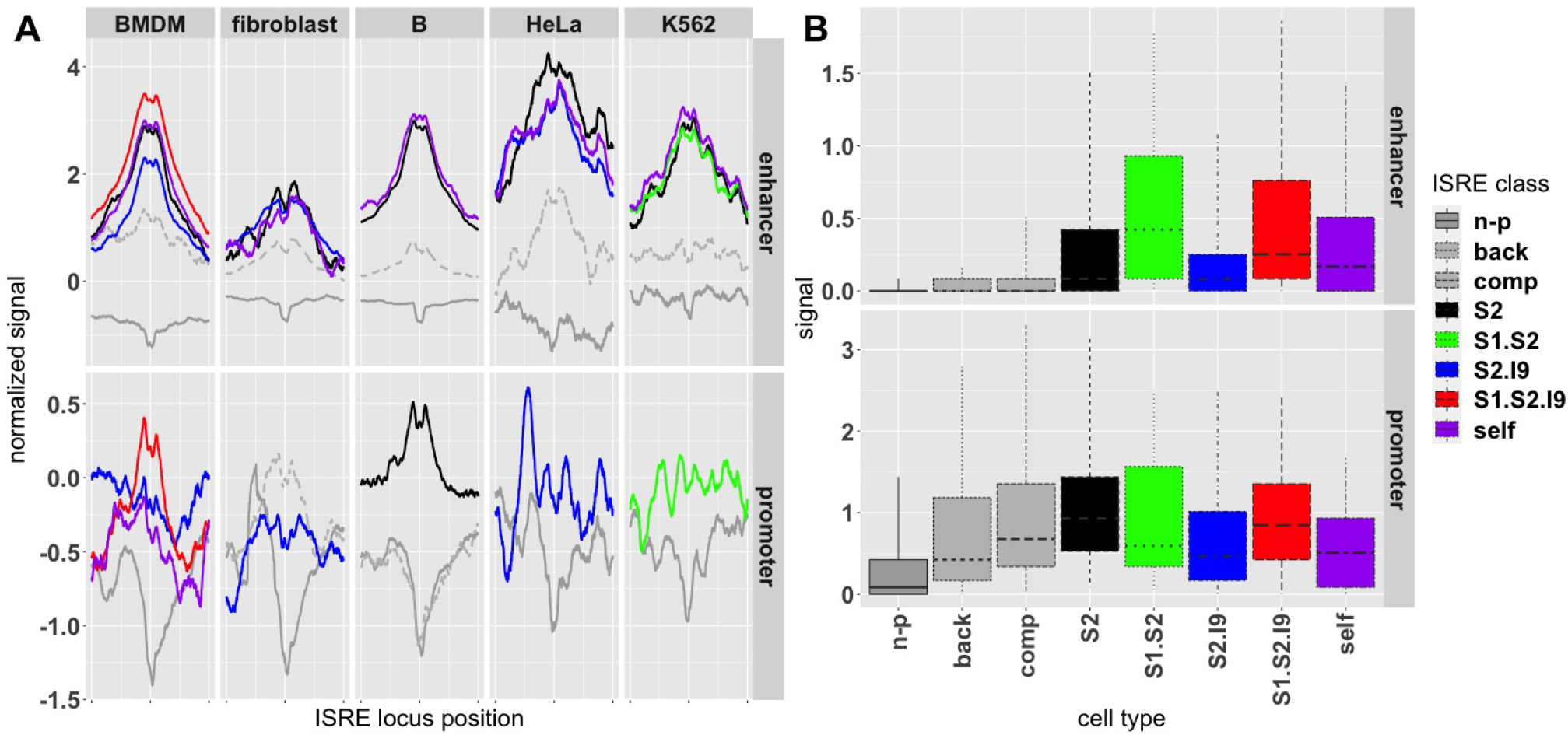
Profiles and Distribution of ATACseq Signals. (A) The mean ATACseq signal over a 1010 base pair locus surrounding each ISRE, split by ISRE class, cell type, and enhancer (top panels) and promoter (bottom panels) regions. The mean signals are normalized so that the background mean signal is 0 at all positions. (B) The distribution of the ATACseq signals in BMDM at the middle position of the 1010 base pair locus. Lower and upper whiskers give 5% and 95% quantiles, respectively, and lower and upper hinges give 25% and 75% quantiles. Signals are shown only for ISRE classes with more than 75 ISREs. Unbound ISRE classes are colored grey. In panel A, background signal is not shown, non-peak signal is solid, and complement signal is dashed. (n-p=non-peak, back=background, comp=complement)

For ISRE in the enhancer region, non-peak ISRE had mean signals that were negative, meaning that ISRE unbound across all cell types were on average less accessible than background, a phenomenon noted by other authors [3, 15]. The bound ISRE classes had positive, mean signals, meaning that bound ISRE had increased accessibility relative to background. ISRE in the complement class had an accessibility level just above background, while ISRE in the self class had higher accessibility levels. Recall, ISRE in the self class were only bound in the given cell type while ISRE in the complement class were only bound in other cell types. Since all ISREs in self and complement classes are bound in some cell type, their differential accessibility reflects cell-specific mechanisms. Across all cell types, bound ISRE in the enhancer had a maximal plateau centered at the ISRE motif and extending roughly 200 base pairs, a profile consistent with the absence of a nucleosome [10].

Mean signals for ISRE in promoter regions were different. In contrast to ISRE in the enhancer region, the bound ISRE class signals were not separated from background. Bound ISRE in the promoter region did not generally exhibit the plateau profile seen in enhancer regions, possibly because nucleosome depletion is common in promoter regions. Non-peak ISRE in promoter regions did show a dip in signal at the ISRE motif position across all cell types, suggestive of nucleosome positioning at the ISRE.

To visualize the distribution of the signals, beyond mean values, we considered the central position of the ATACseq signal in the ISRE loci, position 505 of 1010. Figure 6B shows the distribution of signals at this position for BMDM, again split by enhancer (upper panel) and promoter regions (lower panel), but in this case we did not normalize the signal against background; other cell types had similar distributions for their mean signals. For ISRE in the enhancer region, signals for the bound ISRE classes were elevated above background, while for ISRE in the promoter regions, bound ISRE signals overlapped with background. As might be expected, accessibility was higher for ISRE in the promoter region regardless of the class, reflecting the distinct chromatin state of ISRE between the two regions.

To build classifiers using the homeostatic ATACseq signals, we defined a set of 12 features (values) composed of the mean of the signal across the 1010 base pair ISRE locus and the fraction of the signal contained in eleven, 90 base pair windows covering the locus (the windows did not include the 10 base pairs on the far ends). We built binary classifiers, considering one of the bound ISRE classes against one of the unbound (non-peak, background, or complement) classes or the STAT2 class.

In line with the signal means, bound ISRE in the enhancer region were relatively accurately distinguished from the non-peak class, with an AUC varying from 0.70 to 0.90 depending on the bound ISRE class and precision greater than 0.90 across all bound ISRE classes, see Figure 7A for AUC values and Supplementary Figure S1A for precision values. Prediction in the promoter region was less accurate, with AUC and precision values lower than enhancer values by 0.05 − 0.15 and 0.10 − 0.30, respectively. Nevertheless, bound ISRE could be distinguished from non-peak ISRE in the promoter region with a significant level of accuracy.

**Figure 7:**
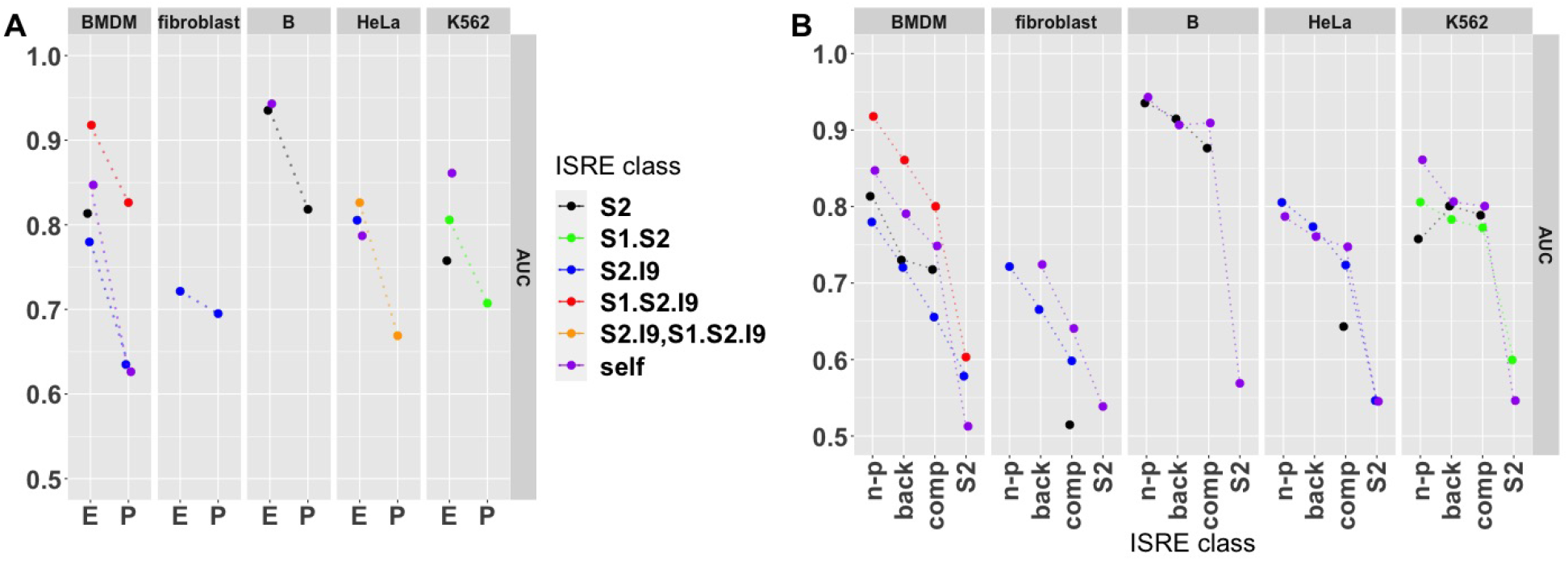
Prediction Accuracy Using ATACseq. (A) AUC for prediction of bound ISRE classes against the non-peak ISRE class for ISRE in the enhancer (E) and promoter (P) regions. (B) AUC of bound ISRE classes against non-peak (n-p), background (back), complement (comp), and STAT2 (S2) classes for ISRE in the enhancer region. Prediction involving ISRE classes with less than 20 ISRE were not considered. Due to limited sample sizes, for HeLa cells we combined STAT2.IRF9 and STAT1.STAT2.IRF ISREs into a single ISRE class (S2.I9,S1.S2.I9)

We next considered prediction accuracy of bound ISRE classes against the unbound ISRE classes and STAT2 ISRE. Promoter regions had less bound ISRE, leading to limitation in our statistical power, so we limited our analysis to enhancer regions, see Methods for details. AUC and prediction accuracy fell sequentially as we compared bound ISRE classes against non-peak, background, complement, and STAT ISRE classes, respectively; see Figure 7B for AUC values and Supplementary Figure S1B for precision values. Across all cell types and bound ISRE classes, AUC and precision against the STAT2 ISRE class were 0.60 or below, a level of accuracy only slightly above the 0.55 cutoff for statistical significance. Chromatin accessibility is therefore not the distinguishing characteristic between the strength in ChIP signal across STAT1.STAT2.IRF9, STAT2.IRF9 ISRE and STAT2 ISRE (recall Figure 5). Complement ISRE occupied a middle ground between non-peak ISRE and background loci, which were accurately distinguished from bound ISRE, and the STAT2 ISRE class. The intermediate level of accuracy could reflect mistaken classification of bound ISRE as complement ISRE, or complement ISRE may simply have a range of chromatin accessibility.

‘

#### Histone Modifications

We next considered signals for H3K4ME1, H3K4ME3, and H3K27AC ChIP datasets, splitting as before by cell type, ISRE class, and enhancer and promoter regions. Figure 8 shows the mean signals for ISRE in the enhancer region. Across all three histone modifications and all cell types, the non-peak ISRE class had reduced signal relative to the background signal, as was the case for ATACseq mean signals, pointing to a general quiescence of ISRE that are not bound. H3K4ME1 and H3K27AC signals for bound ISRE classes were higher than the background class in all cell types. In contrast, H3K4ME3 signals of the bound ISRE classes did not differ from the background signal in BMDM, fibroblast, and K562, but were elevated in HeLa, B, and THP1 cells. As was the case for ATACseq signals, the complement class signal was only slightly increased relative to background while the self class had higher signal than background, again showing cell-specific differences in the homeostatic state. Histone modification signals in the promoter region, differed from those in the enhancer region, see Supplementary Figure S2. The mean background signal in the promoter region did not clearly separate from the mean signal of bound ISREs.

**Figure 8:**
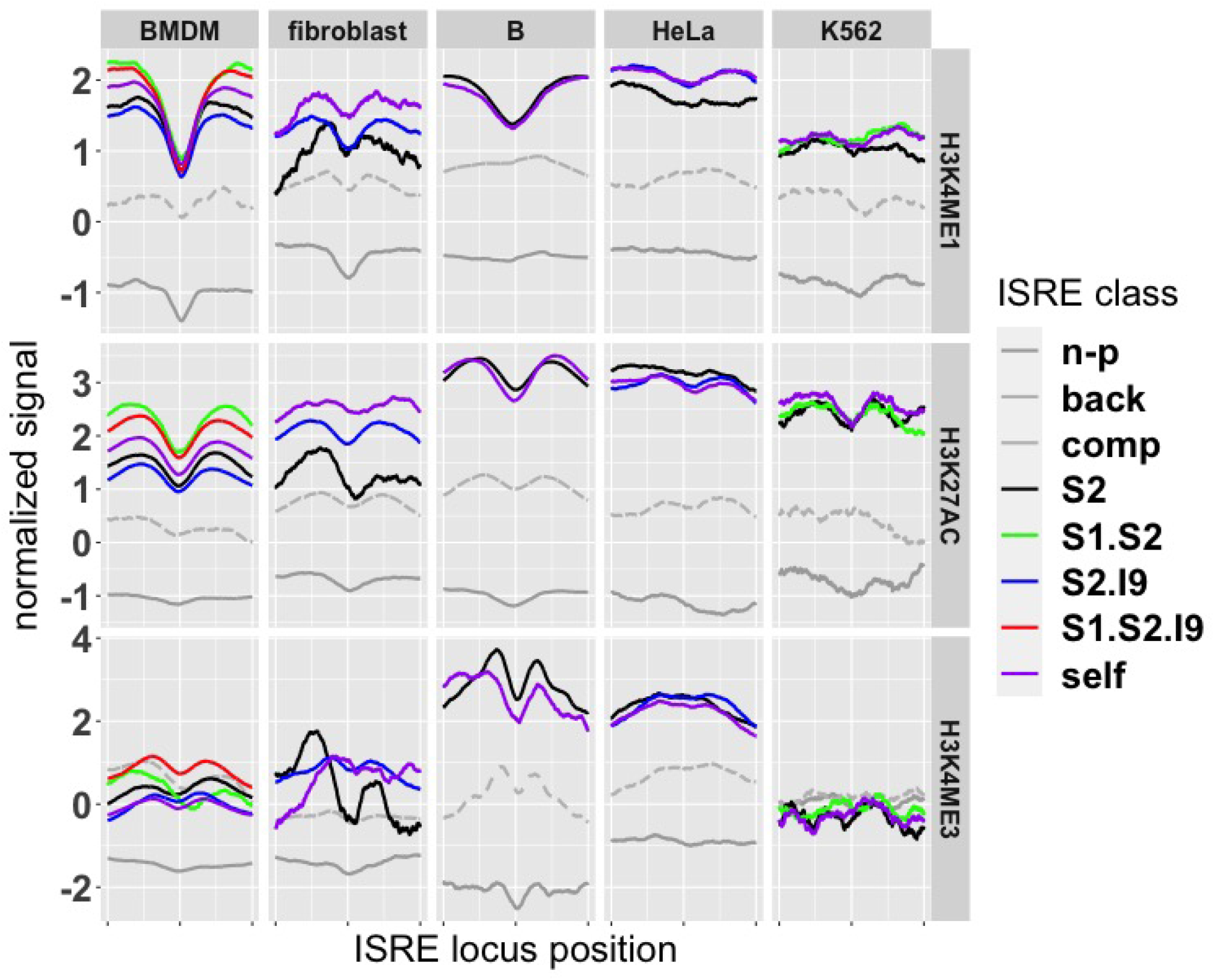
Profiles of Histone Modification ChIPseq Signals. Shown are mean ChIPseq signals, split by cell type and ISRE class, for ISRE in enhancer regions. Bound ISRE signals were separated from background in all cell types for H3K4ME1 and H3K27AC modifications, but not for H3K4ME3 modifications. Across all cell types and histone modifications, non-peak ISRE had lower signals than background. Background signal was normalized to 0 and is not shown, non-peak signal is solid, and complement signal is dashed.

As we did for the ATACseq signal, we built classifiers to predict ISRE class based on the homeostatic, histone signals. We compared classification accuracy using each of the three histone modification ChIPseq signals and using ATACseq signals, see Figure 9. Overall, H3K4ME1 signals had the best predictive accuracy. In contrast, classification using H3K4ME3 had significantly lower prediction accuracy than ATACseq in all cell types but HeLa cells. As was the case for prediction using ATACseq, prediction accuracy fell sequentially as bound ISRE classes were compared against non-peak ISRE, background, complement ISRE, and STAT2 ISRE and, further, prediction accuracy was higher for ISRE in the enhancer region than in the promoter region.

**Figure 9:**
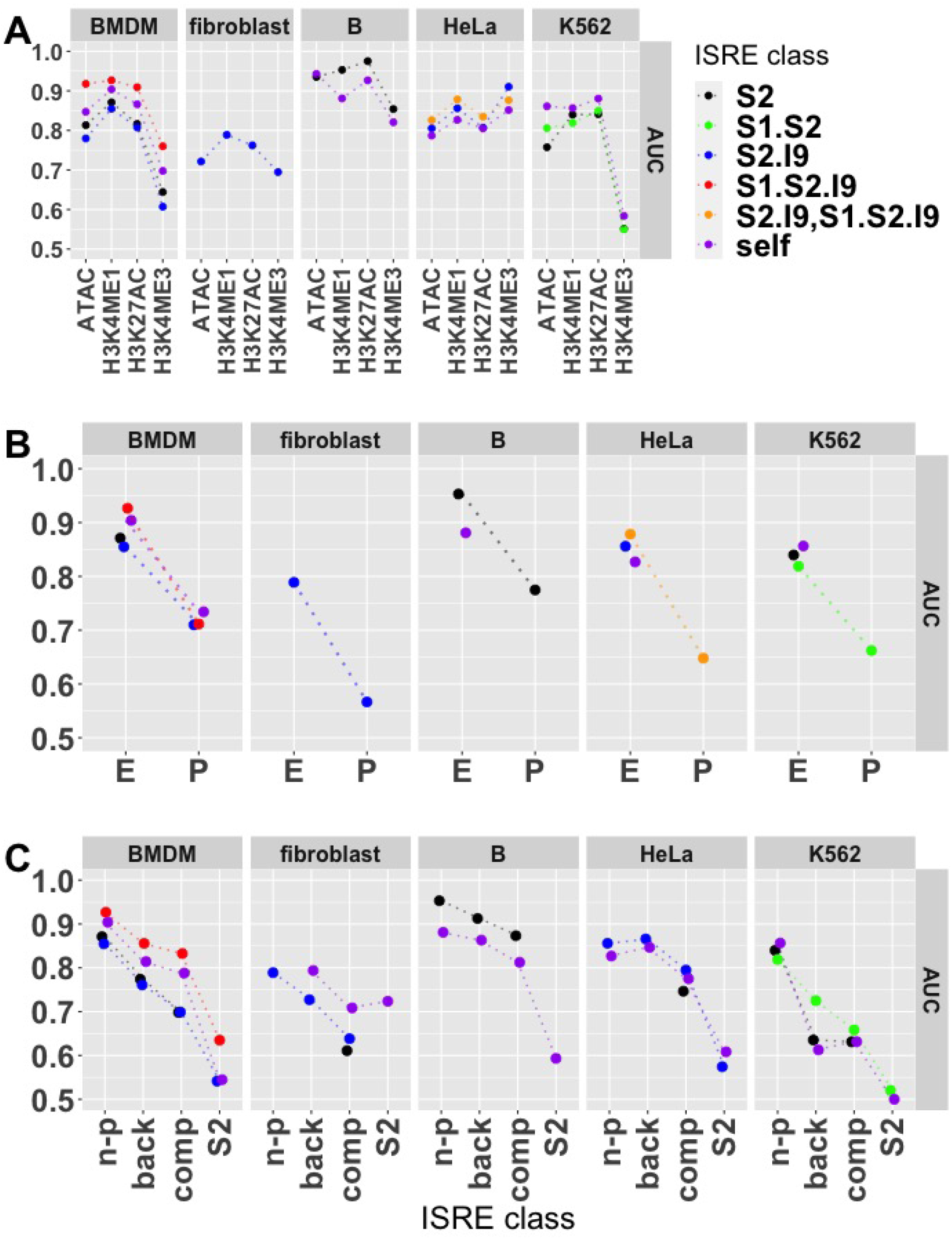
Prediction Accuracy of ISRE Classes Using Histone Modification ChIPseq Signals. (A) AUC for prediction of bound ISRE classes in the enhancer region against non-peak ISRE class using ATACseq and histone modification signals. Overall, H3K4ME1 provided the best accuracy. (B) Comparison of AUC for prediction of bound ISRE classes against the non-peak class in the enhancer (E) and promoter (P) regions using the H3K4ME1 ChIPseq signals. (C) AUC for prediction of bound ISRE in the enhancer region against the different unbound ISRE classes and the STAT2 class using the H3K4ME1 ChIPseq signals. Note the similarity of panels (B) and (C) to results using the ATACseq signal, shown in Figure 7

#### Transcription Factor Motifs

We next considered motifs of different transcription factors (TF) within the 1010 base pair ISRE loci. We considered the 746 TF motifs in the JASPAR vertebrates dataset [27]. For each TF motif and each ISRE locus, we used MEME-FIMO ([31]) to compute a score, reflecting the best possible match of the TF motif over the locus sequence. In scoring TF motifs, we ignored matches involving the 10 base pairs of the ISRE motif on which the locus was centered. Using the 746 scores for each locus as features, we built binary classifiers as we did for the ATACseq and histone, ChIP datasets.

For ISRE in the enhancer region, classification using motifs was not as accurate as classification using H3K4ME1 signals, except for fibroblasts. In contrast, in promoter regions, AUC values using motifs were roughly equal or higher than H3K4ME1 AUC values in all cell types except B cells. See Supplementary Figure S3.

The classifiers selected 100s of motifs as predictive of peak binding. Since our AUC and precision values were computed on a test dataset and comparison between training and test AUC showed only small differences, the inclusion of multiple motifs in the classifications did not represent overfitting. Instead, the large number of motifs likely reflect the correlation between different motifs.

We sought to identify individual motifs with significant levels of prediction accuracy. Since the sample sizes were larger, we focused on ISRE in enhancer regions. We constructed binary classifiers using each of the 746 TF as the sole predictor and identified TF motifs with AUC and precision values that were significantly elevated at an FDR of .05. For each cell type, we collected the 3 most significant motifs in classification of bound ISRE against non-peak ISRE and of self ISRE against complement ISRE, see Table 8. In classification against non-peak ISRE, the JASPAR STAT1::STAT2 motif, which is an ISRE motif, was present in all cell types except HeLa cells, in-line with previous results suggesting multimeric ISRE binding [78, 44]. (Recall, we removed the ISRE motifs on which each ISRE locus was centered in computing motif scores, so the STAT1:STAT2 motif was matching other ISREs within the loci.) Further, we found multiple IRF motifs in BMDM, fibroblast, and K562 cells. In contrast, when we compared self ISRE against complement ISRE, which presumably should be differentiated by cell specific motifs, the STAT1:STAT2 motif and IRF motifs were not present in any cell type and no motifs were shared across the cell types, supporting cell-specific regulation of ISRE loci.

**Table 8:** Single Motifs with Significant Predictive Capacity. We identified motifs that classified bound ISRE against non-peak ISRE (bound vs non-peak) and self ISRE against complement ISRE (self vs complement) with statistically significant AUC and precision. Shown for each cell type are the 3 significant motifs with the highest precision. Motifs that distinguished bound ISRE from non-peak ISRE were enriched for STAT and IRF motifs and were not cell type specific. Motifs distinguishing self ISRE (ISRE bound only in the given cell type) from complement ISRE (ISRE bound in other cell types than the given cell type) were cell type specific.

| cell type | bound vs. non-peak | self vs. complement |
| --- | --- | --- |
| BMDM | STAT1::STAT2,IRF8,IRF9 | ETV6,EHF,FOXF1 |
| fibroblast | IRF1,IRF8,STAT1::STAT2 | HOXA10,HOXA9,FOXD3 |
| B | STAT1::STAT2,ELF3,EHF | SMAD2::SMAD3,LMX1B,DLX2 |
| HeLa | HOXD11,EGR2,SHOX2 | FOSL1::JUN,BATF::JUN,FOSL1::JUNB |

#### Sequence

We next examined whether sequence specificities, not associated with co-transcription factors, were predictive of ISRE class. To encode the nucleotide sequence of the ISRE motif at the center of the locus, we considered a 30 base pair window starting 10 base pairs upstream of the ISRE motif and ending 10 base pairs downstream. For each position within the window, we defined features specifying the position’s nucleotide, an encoding similar to a position weight matrix (PWM) of a motif in that positions are considered independently, so we refer to these features as PWM features.

We also considered DNA shape over the same 30 base pair window. Mathelier et al. calculated DNA shape features from DNA sequence [49] and we used the R package DNAshapeR, [12], to compute 5 different shape features (minor groove width, roll, propeller twist, helix twist, and electrostatic potential) using a five base pair window centered at each base pair in our 30 base pair window, giving us 150 shape features. Finally, we also considered CpG content in the 1010 base pair locus containing each ISRE. To define CpG features, we defined as features the total number of CpG pairs in the locus as well as the fraction of CpG pairs within 11 windows decomposing the locus (as we did for the ATACseq and histone signals).

We built binary classifiers using CpG, PWM, and shape features as well as classifiers that combined all these features. For ISRE in the enhancer region, the PWM, and shape features had roughly the same AUC and precision, in the range 0.7 0 *−*.90 and 0.80 − 0.95 respectively, while CpG features had low AUC and precision, see Figure 10A. Combining PWM and shape features raised AUC by up to 0.10. Across all cell types but BMDM, accuracy for ISRE in the promoter region were similar to AUC for ISRE in the enhancer which was notably not the case using ATACseq and histone ChIPseq signals. As before, we classified bound ISRE against non-peak, complement, and STAT ISRE classes. (In this case, we did not include the background class because the lack of an ISRE motif meant that background loci were completely distinguished from bound ISRE.) In contrast to ATACseq and histone signals, classification of bound ISRE against the complement class had AUC substantially less than AUC against the non-peak class across all cell types, suggesting that the ISRE sequence is a strong predictor of ISRE that are never bound, but a weak predictor of cell-specific ISRE binding; see Figure 10B.

**Figure 10:**
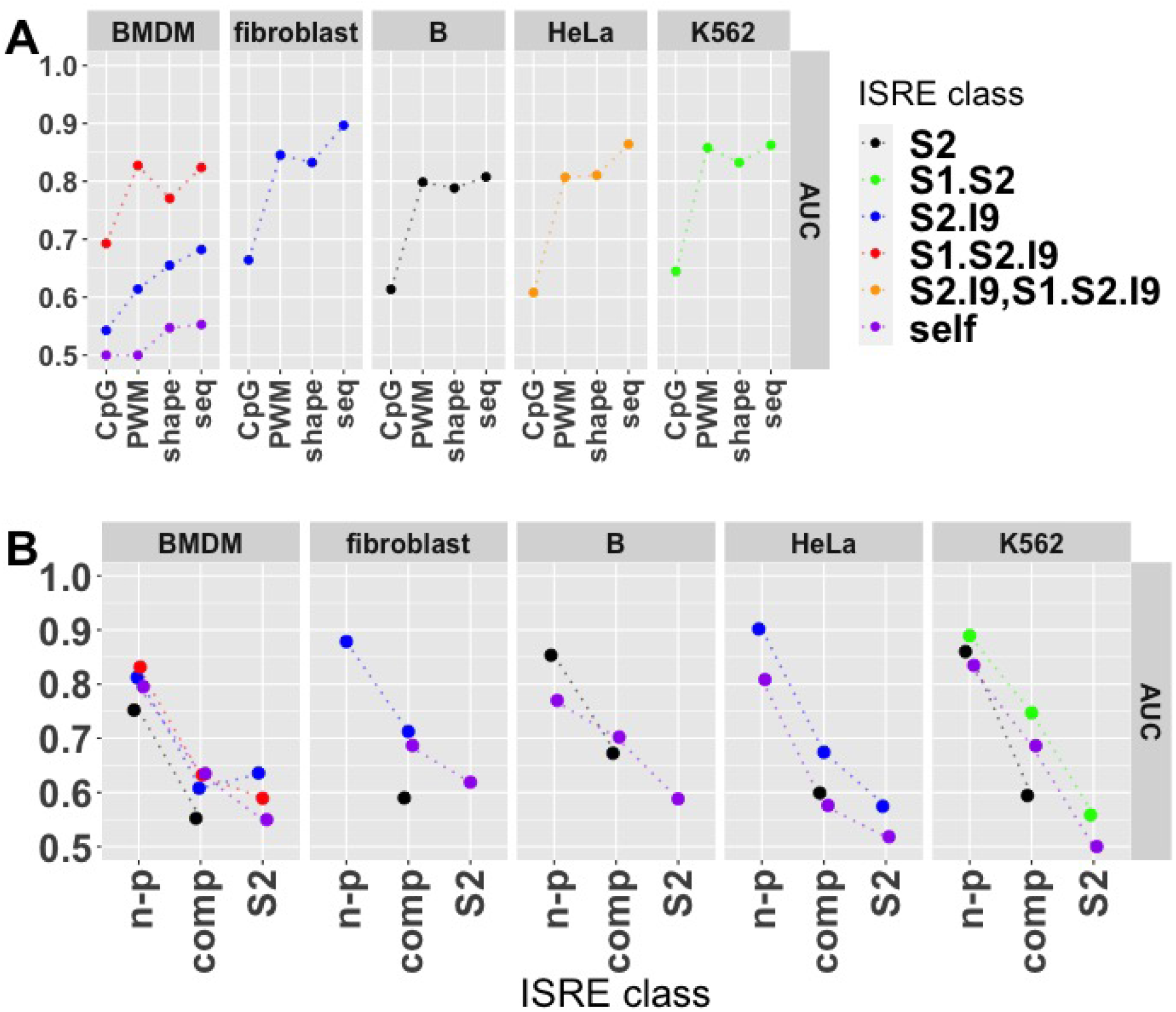
Prediction Accuracy Using DNA Sequences. AUC for prediction of bound ISRE classes in the enhancer region against (A) the non-peak ISRE class using CpG, PWM, shape feature types and all three of these feature types combined (seq), and (B) against the different unbound ISRE classes and the STAT2 class using the combined features (seq).

To gain a better understanding of sequence patterns that were predictive of ISRE class, we considered the contribution of the different nucleotides at each position in the 30 base pair window to the probability an ISRE was bound according to our classifiers. Figure 11 shows the relative importance of nucleotides in distinguishing STAT2.IRF9 and STAT1.STAT2.IRF9 ISRE from background, non-peak, complement and STAT2 ISRE in the enhancer region of BMDM; other cell types showed similar results. Positive and negative values are nucleotides that increased and decreased, respectively, the probability an ISRE was in the STAT2.IRF9 or STAT1.STAT2.IRF9 classes. When STAT2.IRF9 or STAT1.STAT2.IRF9 ISRE were compared to background, we saw the canonical TTTCNNTTTC pattern, reflecting the absence of an ISRE motif on background loci. In contrast, in comparison to non-peak ISRE, the STAT2.IRF9 or STAT1.STAT2.IRF9 ISRE were enriched in the canonical TTTC pattern at the 5’ end of the motif and a TT pattern in the 3’ end, but each position is less predictive than in the case of background. We do not see the typical TTTCNNTTTC motif because all the ISRE in the non-peak class are within one nucleotide of this motif. For comparison of STAT2.IRF9 and STAT1.STAT2.IRF9 ISRE to STAT2 ISRE, there was enrichment for *G* at the 2nd position in the ISRE motif, suggesting STAT2.IRF9 and STAT1.STAT2.IRF9 ISREs were enriched for the TGTTNNTTTC motif relative to STAT2 ISREs, but all other positions had weak or no contribution to differentiating the classes. Finally, there was essentially no enrichment in comparing the STAT2.IRF9 and STAT1.STAT2.IRF9 ISRE to complement ISRE, suggesting that cell specificity of ISRE binding is not mediated through ISRE sequence.

**Figure 11:**
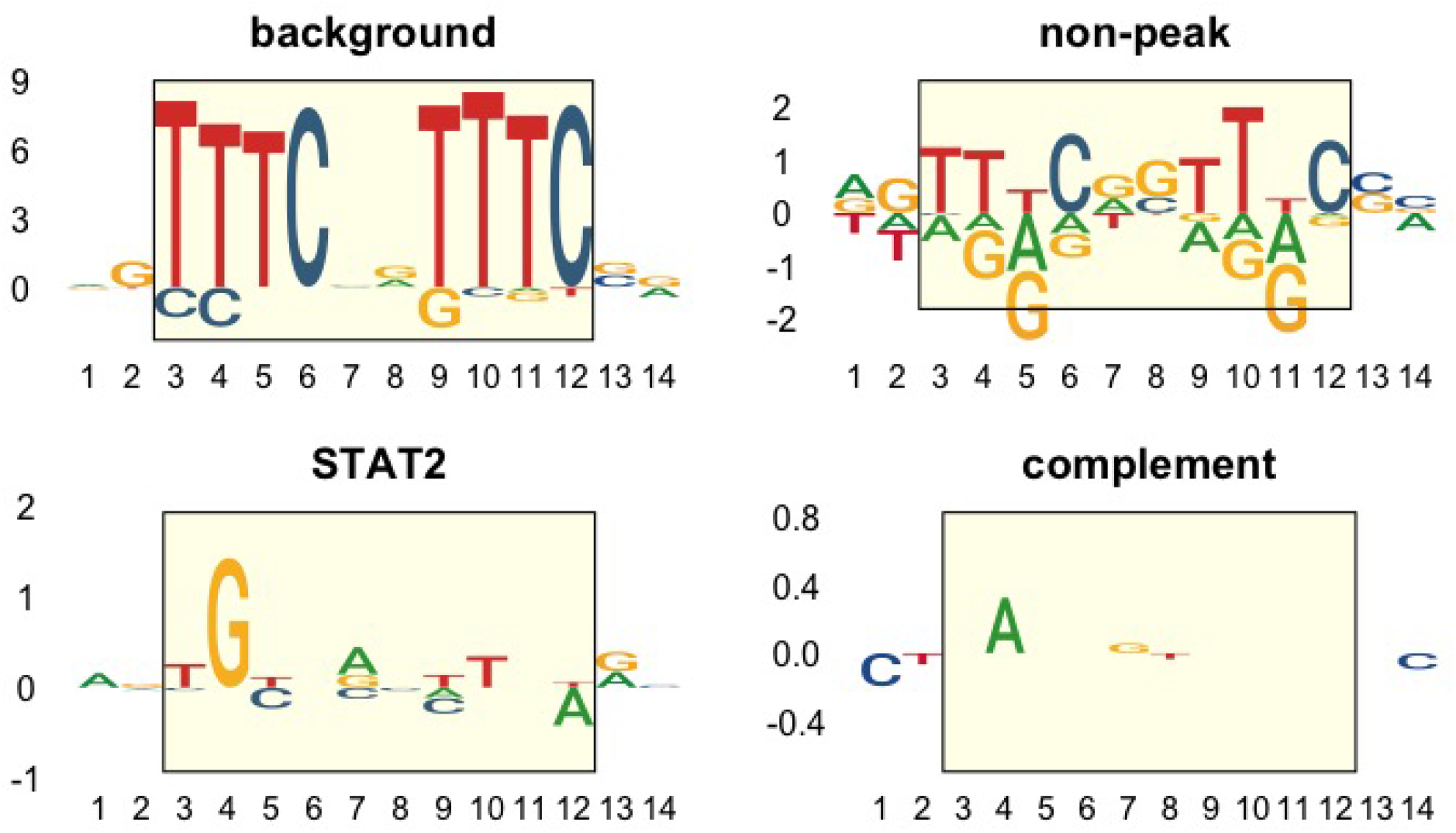
Visualization of Predictive Capacity of PWM Features. For each position in a 30 base pair window including the 10 base pair ISRE motif, we used the presence of a A,C,G, and T as a feature to classify STAT2.IRF9 and STAT1.STAT2.IRF9 ISRE classes against background, non-peak, complement, and STAT2 ISRE. Shown are the contributions of the different nucleotides at the different positions to the probability that an ISRE is classified in the STAT2.IRF9 and STAT1.STAT2.IRF9 classes. Positive and negative values mean that the nucleotide increased and decreased, respectively, the probability. Shown are the 10 base pairs covering the ISRE motif (yellow box) and the 2 base pair positions up and downstream.

#### Homeostatic STAT1, STAT2, and IRF9 signals

We next considered the homeostatic signals of STAT1, STAT2, and IRF9 ChIP datasets, in a manner completely analogous to our treatment of ATACseq and histone ChIP signals. For K562 cells, we did not have STAT1 and STAT2 ChIPseq data at homeostasis. Instead of homeostasis, K562 STAT1 and STAT2 ChIPseq datasets were collected at 30 minutes post IFN stimulation, as opposed to the IFN stimulated datasets for K562 cells, which were collected at 6 hours.

Figure 12 shows the homeostatic, mean STAT1, STAT2, and IRF9 signals. In contrast to results for ATACseq and histone signals, mean signals of bound ISRE in enhancer and promoter regions were similar once normalized against background mean signals. The mean signals of the non-peak class were essentially zero, even for K562 cells, in-line with the expectation that background loci and non-peak ISRE would both have essentially no STAT1, STAT2, or IRF9 binding. The bound ISRE classes had positive, mean signals in all cell types and both promoter and enhancer regions, except for the enhancer region of fibroblasts. K562 signals were particularly elevated, reflecting the 30 minutes of IFN stimulation. ISRE in the complement class had a small positive or zero, mean signal, while ISRE in the self class had elevated mean signal across cell types, supporting the presence of cell specific binding at homeostasis.

**Figure 12:**
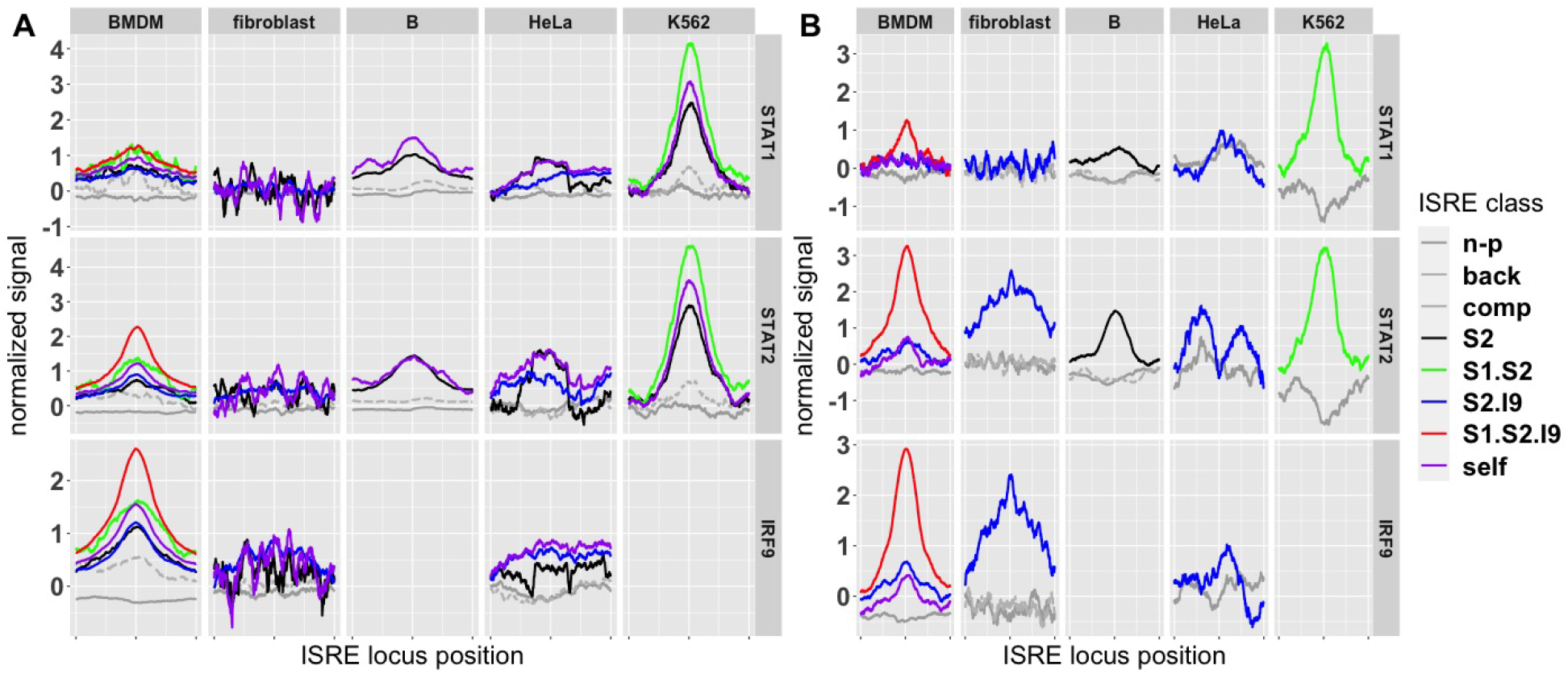
Profiles of Homeostatic STAT1, STAT2, and IRF9 ChIPseq Signals. Shown are mean ChIPseq signals, split by cell type and ISRE class, for ISRE in (A) enhancer regions and (B) promoter regions. All signals are from datasets collected at homeostasis, except for K562 cells, which were sampled 30 minutes after IFN stimulation. Background signal was normalized to 0 and is not shown; non-peak signal is solid and complement signal is dashed.

Before turning to binary classifiers, we considered calling ChIP peaks as we did for the ChIP datasets collected under IFN stimulation. We called peaks, associated peaks to ISRE, and split bound ISRE into classes exactly as we did for datasets under IFN stimulation. Ignoring K562 cells, across all cell types there were relatively few ISRE bound at homeostasis, at least as called by MACS2 using an FDR of 0.01. The number of ISRE bound at homeostasis was between 1% − 5% of the number bound under IFN stimulation for all cell types, except THP1, for which the number at homeostasis was 17% of the number under IFN stimulation. Across all cell types less than 6% of ISRE bound under IFN stimulation were also bound at homeostatic conditions. Conversely, a substantial percentage of ISRE bound at homeostasis were also bound under IFN stimulation. For BMDM and splenic B cells, 96% and 66% of ISRE bound at homeostatic were bound under IFN stimulation. For fibroblast, HeLa, and THP1, the percentage varied between 16% − 36%. See Table 9 for details.

**Table 9:** Homeostatic Bound ISRE.. Just as we identified bound ISRE under IFN stimulation, we identified bound ISRE at homeostasis. Shown are the number of bound ISRE under IFN stimulation (IFN) and at homeostasis (homeo), the number of ISRE that were bound under IFN stimulation and homeostasis (joint), the fraction of bound ISRE under IFN stimulation that were also bound at homeostasis (joint (IFN)), and the fraction of bound ISRE at homeostasis that were also bound under IFN stimulation (joint (homeo)). K562 peaks reflect ChIPseq datasets collected at 30 minutes after IFN stimulation rather than at homeostasis.

| cell type | IFN | homeo | joint | joint (IFN) | joint (homeo) |
| --- | --- | --- | --- | --- | --- |
| BMDM | 11818 | 496 | 476 | 0.04 | 0.96 |
| fibroblast | 3287 | 44 | 7 | 0.00 | 0.16 |
| B | 2760 | 142 | 36 | 0.01 | 0.25 |
| HeLa | 890 | 12 | 8 | 0.01 | 0.67 |
| K562 | 874 | 308 | 176 | 0.20 | 0.57 |
| THP1 | 509 | 85 | 31 | 0.06 | 0.36 |

As we did for previous homeostatic factors, we built classifiers to predict ISRE class using homeostatic, STAT1, STAT2, and IRF9 signals individually, and jointly. We refer to the joint features of the STAT1, STAT2, and IRF9 signals as ISGF3 features. Prediction accuracy using IRF9 or ISGF3 features were roughly equivalent, at least for the cell types for which we had IRF9 ChIPseq datasets. IRF9 was a better predictor than STAT2, which was a better predictor than STAT1. For ISGF3 features, AUC was better than 0.80 in BMDM and B cells, but less than 0.70 in HeLa and fibroblast cells. We speculate that the difference reflects greater ISRE binding at homeostasis in immune cells than non-immune cells. Interestingly, and somewhat reassuringly, AUC values for K562 cells were greater than 0.95, showing that STAT1 and STAT2 binding at 30 minutes post IFN stimulation accurately predicts ISRE binding at 6 hours. As opposed to prediction accuracy using ATACseq or histone signals, accuracy for ISRE in promoter regions were either roughly equal or better than in the enhancer region, see Figure 13.

**Figure 13:**
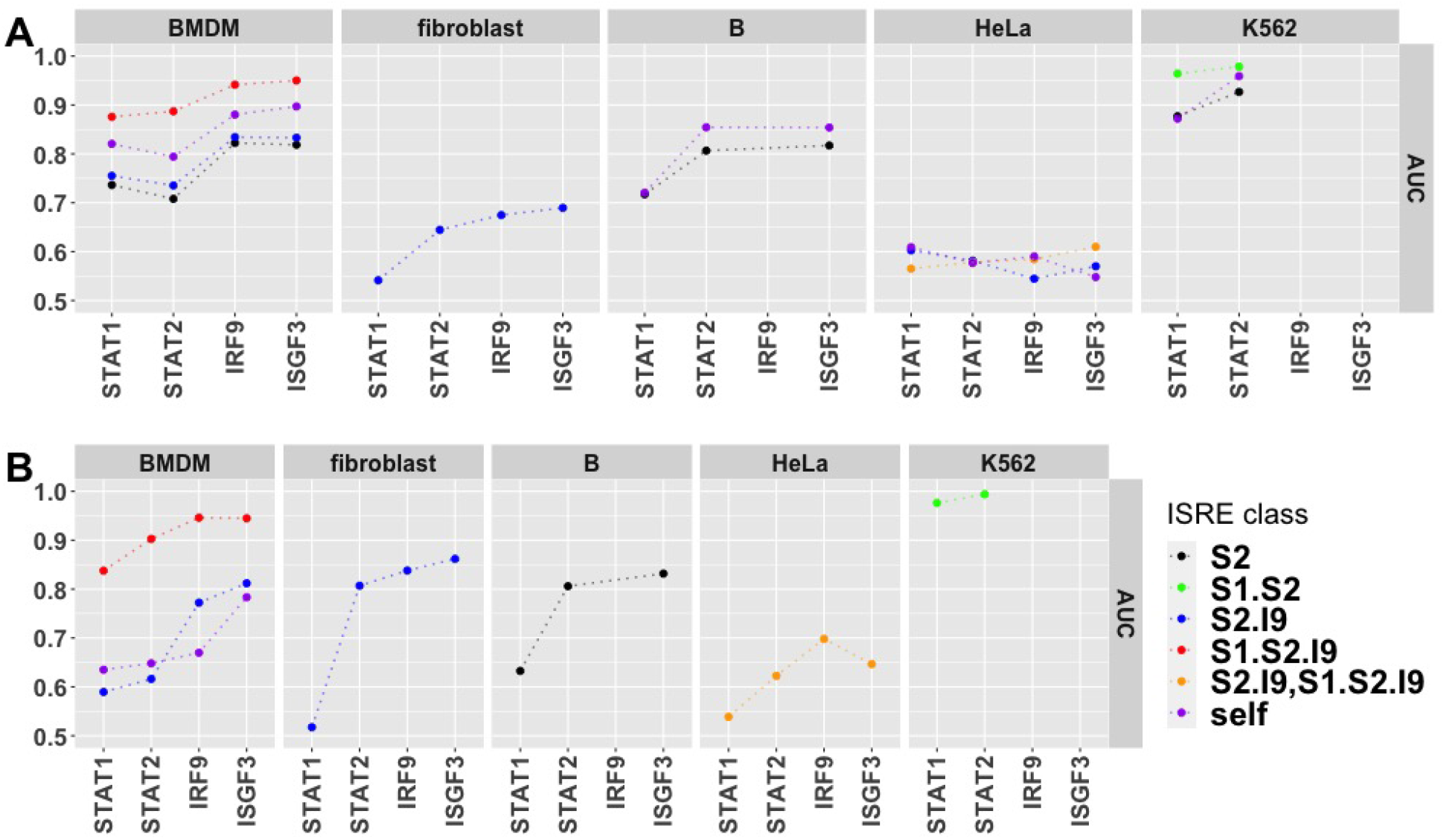
Prediction Accuracy Using STAT1, STAT2, and IRF9 ChIPseq Signals. AUC for prediction of bound ISRE against non-peak ISRE in the enhancer (A) and promoter (B) regions.

#### Joint Prediction

We next compared prediction accuracy associated with combinations of our feature types: chromatin features, formed by combining the ATACseq and histone features, sequence features, formed by combining PWM and shape features, ISGF3 features, formed by combining STAT1, STAT2, and IRF9 features, and motif features. For ISRE in enhancer regions, prediction accuracy increased sequentially as we used motif, sequence, and chromatin features respectively, except in fibroblast while for ISRE in the promoter regions, sequence features had the highest accuracy across fibroblast, HeLa, and K562, and were only slightly below chromatin features in BMDM and B cells. Prediction accuracy for ISGF3 features were relatively high in some cell types and low in others; overall sequence and chromatin features were better predictors; see Figure 14.

**Figure 14:**
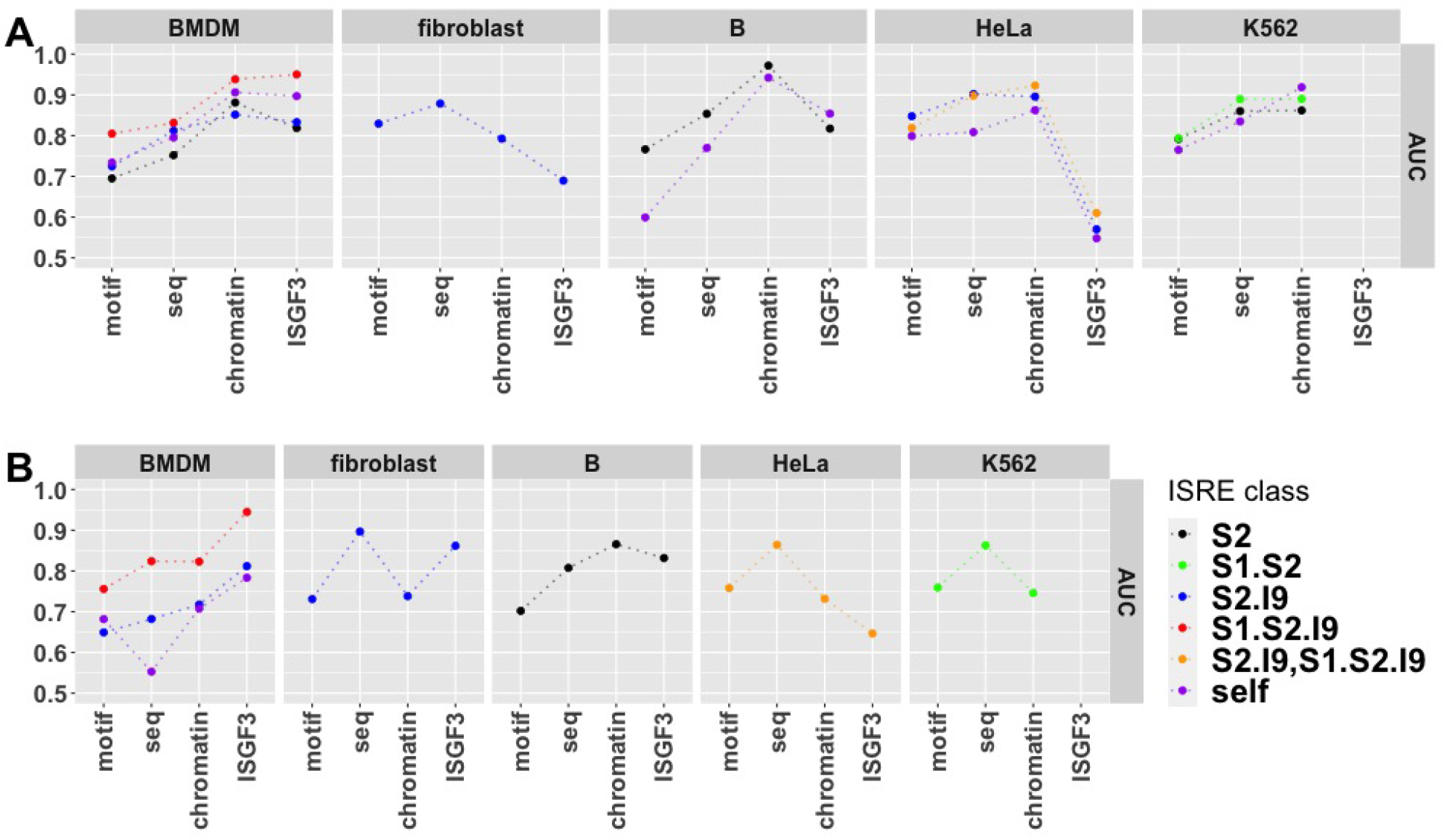
Prediction Accuracy Using Different Feature Types. AUC for prediction of bound ISRE against non-peak ISRE in the enhancer (A) and promoter (B) regions.

To understand the full prediction accuracy of the homeostatic state. we considered prediction accuracy using chromatin features, chromatin features combined with sequence features, and the chromatin, sequence, and ISGF3 features combined; see Figure 15. Motif features made little difference in combination with the other feature types. In line with previous results [85], adding sequence features to chromatin features raised AUC by 0.05 − 0.10, although in fibroblast the difference was greater, 0.15. Adding ISGF3 features to chromatin and sequence features raised AUC by a further 0.05 − 0.10. Notably, when we used all homeostatic features, we could distinguish a bound ISRE from a non-peak ISRE in the enhancer region with AUC greater than 0.90 in all cases. For promoter regions, there was a similar increasing trend, but AUC values were roughly 0.10 lower than AUC values for enhancer regions. Finally, when we used all homeostatic features to classify bound ISRE against non-peak, complement, and STAT2 ISRE classes, we found that AUCs dropped sequentially. AUC values for classification of bound ISRE against complement ISRE were between 0.75 − 0.90, demonstrating that ISRE are associated with cell-specific homeostatic states. AUC values for classification of bound ISRE against STAT2 ISRE were less than 0.70 suggesting that STAT2 ISRE were misclassified or that our homeostatic features or the homeostatic state generally do not include factors that distinguish ISRE bound by STAT2 from ISRE bound by other ISGF3 component combinations.

**Figure 15:**
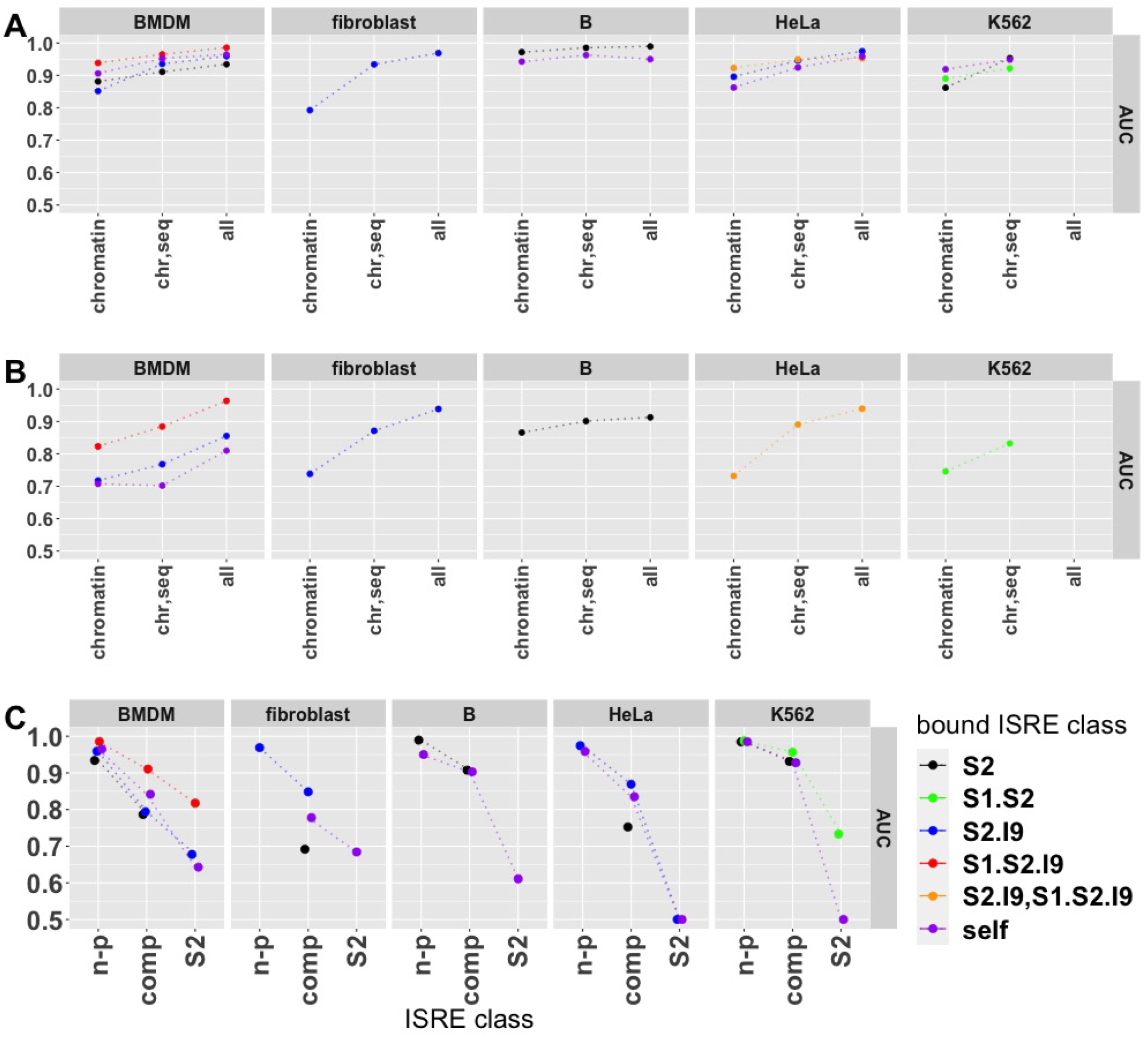
Prediction Accuracy Using All Features. AUC for prediction of bound ISRE against non-peak ISRE in the enhancer (A) and promoter (B) regions and (C) against non-peak, complement and STAT2 ISRE in the enhancer region.

## 4 Discussion

In this work, we examined ISRE binding signatures under type I IFN stimulation in six cell types. Within each cell type, bound ISREs were most dense in promoter regions, but most (70% − 80%) bound ISREs were more than 3kb away from a TSS, in what we referred to as enhancer regions. Most genes (*>*60%) and a substantial percentage of ISGs (*>*25%) with bound ISREs had bound ISREs in their enhancer region. We further found that a substantial percentage of genes and ISGs (*>*40% and *>*10% respectively) with bound ISRE had their bound ISRE restricted to enhancer regions, although this result depends on ChIPseq sensitivity We found that bound ISRE in promoters were shared across cell types at higher levels (41% − 92%) than in enhancers regions (6% − 31%). Using GO ontology enrichment analysis, we found that a significantly greater fraction of bound ISRE in promoters were associated with genes enriched for the viral defense GO class than bound ISRE in enhancers. Overall, these results are consistent with the general paradigm of greater cell specificity in enhancer regulation than promoter regulation.

Presumably, ISREs function to induce or restrict expression of ISGs. In keeping with canonical models of ISG regulation, we found that 25% − 69% of genes with bound ISRE in the promoter were ISGs, even in cases where bound ISRE were restricted to the promoter region In this context, what is perhaps surprising, although other authors have found similar results [32, 74], is that many genes with bound ISRE in the promoter were not ISGs. The role of enhancers in regulating ISGs is more difficult. Comparing genes with no bound ISRE within 100kb of their TSS and genes with a bound ISRE just in the enhancer region, 1% − 3% and 5% − 11%, respectively, were ISGs, a relatively small effect though statistically significant. This small effect could be due to misclassification errors or a functionally weak contribution to differential expression by ISRE in enhancer regions. Alternatively, enhancer regulation may only take place in certain contexts beyond the presence of IFN stimulation, or the timing of differential expression may be delayed beyond the 2 − 6 hour window post stimulation that we consider. Genes with a bound ISRE in the promoter and enhancer regions were more likely to be ISGs than genes with a bound ISRE just in the promoter region, suggesting that bound ISRE in enhancer and promoter regions work synergistically even within the first few hours of IFN stimulation.

We also characterized the ISGF3 components that bound ISRE across cell type. We found that most ISRE binding involved ISGF3 or STAT2:IRF9 dimers, consistent with current models of ISRE binding [25]. However, we also found a substantial percentage (5%-18%) of ISRE associated with only STAT2 peaks. Genes were more likely to be ISGs if bound by ISGF3 than if bound by STAT2:IRF9 dimers and, in turn, were more likely to be ISGs if bound by STAT2:IRF9 than if bound by STAT2. The lower association of ISGs with STAT2:IRF9 binding relative to ISGF3 binding is in-line with previous work showing delayed, differential expression of ISGs under STAT2:IRF9 binding [7]. The low level of differential expression under STAT2 binding of ISRE suggests non-functionality or misclassification. ChIPseq peak summit heights were sequentially greater as we considered STAT2, STAT2:IRF9, and ISGF3 bound ISRE, further supporting a functional hierarchy.

Differences in ISRE signatures across cell types presumably reflect differences in the cell types homeostatic state. Characterizing a cell’s homeostatic state and using that characterization for prediction of transcription factor binding represents a general, open problem in computational biology, and previous work has identified epigenetic state and sequence specificities as predictors of transcription factor binding. e.g. [71, 80, 41, 70]. With this in mind, we split the homeostatic state into different feature types: chromatin features (formed from ATACseq and histone ChIPseq data), motif features (calculated based on transcription factor motifs found in ISRE loci), sequence features (calculated based on the specific DNA sequence forming the ISRE), and ISGF3 features (calculated based on STAT1, STAT2, and IRF9 ChIPseq datasets collected at homeostasis). Importantly, we did not consider transcription levels at homeostatis, although previous work suggests that the IFN response is regulated by homeostatic expression levels, particularly expression levels of STATs [75, 30, 81, 43].

Using these homeostatic features, for each cell type, we built classifiers to predict bound ISRE against unbound ISRE. The most accurate prediction was of bound ISRE against ISRE unbound in all our cell types, which we referred to as non-peak ISRE. In the enhancer region, prediction using chromatin and sequence features had AUC of roughly 0.90 and combination of these two features did better than either individually. In promoter regions, chromatin and sequence features had an AUC of roughly 0.80. Prediction using sequence features was as accurate in promoter regions as enhancer regions, while prediction using chromatin features was not as accurate, possibly because regions near TSS are general accessible and bear histone modifications. We speculate that nucleosome characteristics, which have been shown to regulate ISG expression [39, 1, 40] and which our homeostatic state does not capture, underlie the relatively low prediction accuracy of chromatin features in promoter regions.

We also investigated whether the homeostatic, cell state could predict bound ISRE against ISRE unbound in a given cell type but bound in another cell type, which we referred to as complement ISRE. Intuitively, this prediction is harder than predicting against non-peak ISRE because ISRE that are bound in some cell type should be more similar in state to each other than to ISRE that are never bound. Our analysis of ATACseq signals showed that ISRE bound in a given cell type were more accessible than complement ISRE (ISRE not bound in that cell type but bound in another cell type), supporting a model in which cell-specific transcription factors contribute to opening chromatin around ISRE. Using all features, we found that bound ISRE could be distinguished from complement ISRE at AUC levels of 0.80 0 *−*.90, further supporting cell-specific differences in ISRE homeostatic state.

Finally, we attempted to classify ISRE bound by STAT2:IRF9 dimers or ISGF3 against ISRE bound just by STAT2. Given our results suggesting that STAT2 binding may be non-functional, we reasoned that there may be chromatin differences, for example in accessibility, that restricted peaks to binding by just STAT2. However, prediction of STAT2:IRF9 or ISGF3 ISRE against a background of STAT2 ISRE had low accuracy, suggesting that either the ISRE that we classify as bound by just STAT2 reflect ChIPseq noise, or that other factors not captured by our homeostatic features differentiate ISRE classes.

The main limitation of this work is the dependence on six STAT1, STAT2, and IRF9 ChIPseq datasets, one for each cell type. We used the STAT1, STAT2, and IRF9 ChIPseq datasets collected under IFN stimulation to determine the bound ISRE, which in turn affected every other analysis we did. We cannot rule out that some of our results reflect the bias of a particular ChIPseq dataset. Further, more datasets would allow us to better investigate cell type specific differences. For example, prediction for B cells was particularly accurate, but whether this was associated with the specific dataset or reflective of homeostatic regulation in B cells is unclear. The relative scarcity of STAT1, STAT2, and IRF9 ChIPseq datasets, for example in the GEO and ENCODE databases, has been previously noted [2] and limited our analysis.

Another important limitation involves our measurement of prediction accuracy. Using AUC and precision, we measured our ability to distinguish a typical bound ISRE against, say, a typical non-peak ISRE, rather than attempt to distinguish all bound ISRE from all non-peak ISRE. Indeed, there are millions of unbound ISREs and our prediction accuracy would not separate the full set of unbound ISREs from bound ISREs. On the other hand, we used exclusively linear classifiers, so our prediction accuracy would likely improve if we used non-linear classification, for example a deep learning approach. However, the modest number of bound ISREs might limit our ability to fit non-linear classifiers.

Overall, we have shown that ISRE signatures both overlap and vary between cell types, reflecting cell type specificity in enhancer regions and a conserved antiviral response in promoter regions. We have shown that ISRE signatures correlate with ISG signatures, with bound ISREs in promoters strongly correlated with differential expression of ISGs, but with bound ISREs in enhancers also correlating with differential expression, although more weakly. Finally, we have shown that cellular epigenetics at homeostasis are predictive of ISRE binding under IFN stimulation. Further study may elucidate the homeostatic pathways that shape ISRE signatures and, in turn, ISG signatures.

## Acknowledgments

We are deeply indebted to the many groups that collected the datasets used in this study. In particular, this work would not have been possible without the BMDM, fibroblast, and THP1 data collected by Platanitis et al. in [61] and the HeLa data collected by Au-Yeung and Horvath in [1], and we thank T. Decker and E. Platanitis for answering our questions relating to their datasets. This work benefited from data assembled by the ImmGen consortium, through the B cell data collected by Mostafavi et al.[52], and the ENCODE consortium, through the K562 dataset.

