## Supplementary Tables and Figures for "Characterization and Prediction of ISRE Binding Patterns Across Cell Types Under Type I Interferon Stimulation"

immediate

| cell type | accession | replicates |
|---|---|---|
| BMDM | GSE106702 | 2 |
| | GSM940891, GSM940892 | 2 |
| | GSE66774 | 2 |
| | GSM2974670, GSM2974671 | 2 |
| fibroblast | GSE104529 | 2 |
| | ENCSR000CAZ | 2 |
| | ENCSR000CBA | 2 |
| | ENCSR000CDI | 2 |
| B | GSE75262 | 2 |
| | GSE75250 | 4 |
| | GSM521420 | 1 |
| | GSE51004 | 2 |
| HeLa | GSE106145 | 2 |
| | ENCSR000APW | 2 |
| | ENCSR000AOF | 2 |
| | ENCSR000AOC | 2 |
| K562 | GSE131447 | 3 |
| | ENCSR000EWC | 2 |
| | ENCSR668LDD | 2 |
| | ENCSR000AKP | 2 |
| THP1 | GSM2544216, GSM2544217 | 2 |
| | GSE123872 | 2 |
| | GSE107851 | 3 |
| | GSE107851 | 3 |

Table S1: **Homeostatic, ATACseq and Histone Modfication Datasets.**

| cell type | accession | replicates |
|---|---|---|
| BMDM | GSE115433 | 2 |
| fibroblast | GSE128107 | 1 |
| B | GSE75252,GSE75253,GSE75250,GSE75251 | 4 |
| HeLa | GSE110067 | 1 |
| K562 | ENCSR000FAV,ENCSR000FAU,ENCSR000FAT,ENCSR000FBC | 2 |
| THP1 | GSE128111 | 1 |

Table S2: **STAT1, STAT2, and IRF9 ChIPseq Datasets.**

| cell type | accession | replicates |
|---|---|---|
| BMDM | GSE115434 | 3 |
| fibroblast | GSE128110 | 3 |
| B | GSE75202 | 2 |
| HeLa | GSE15805 | 2 |
| K562 | ENCSR000CWI | 1 |
| THP1 | GSE128113 | 3 |

Table S3: **Transcription Datasets.**

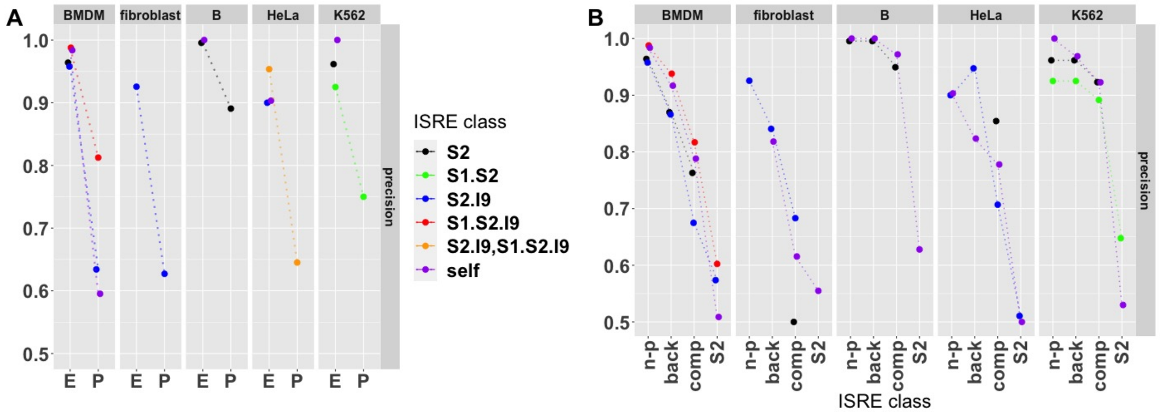


Figure S1: **Precision of Prediction of ISRE Classes Using ATACseq Signals**. (A) Precision, at a recall of 25%, for prediction of bound ISRE classes against the non-peak ISRE class. Prediction involving ISRE classes with less than 20 ISRE were not considered. Precision reflects weighting of samples, see Methods. (B) Precision of bound ISRE classes against non-peak, background, complement, and STAT2 classes.

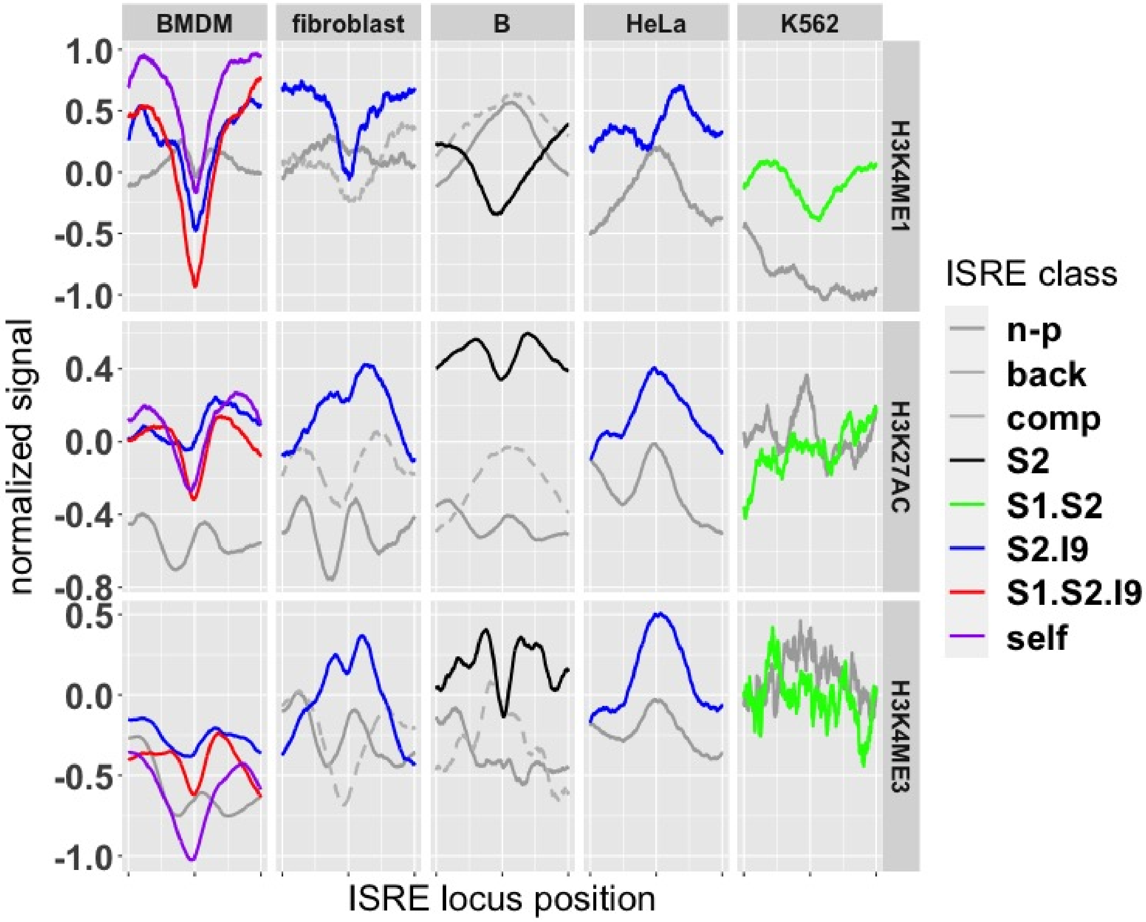


Figure S2: **Profiles of Histone Modification ChIPseq Signals Across ISRE Classes**. Mean signals for ISRE in the promoter region..

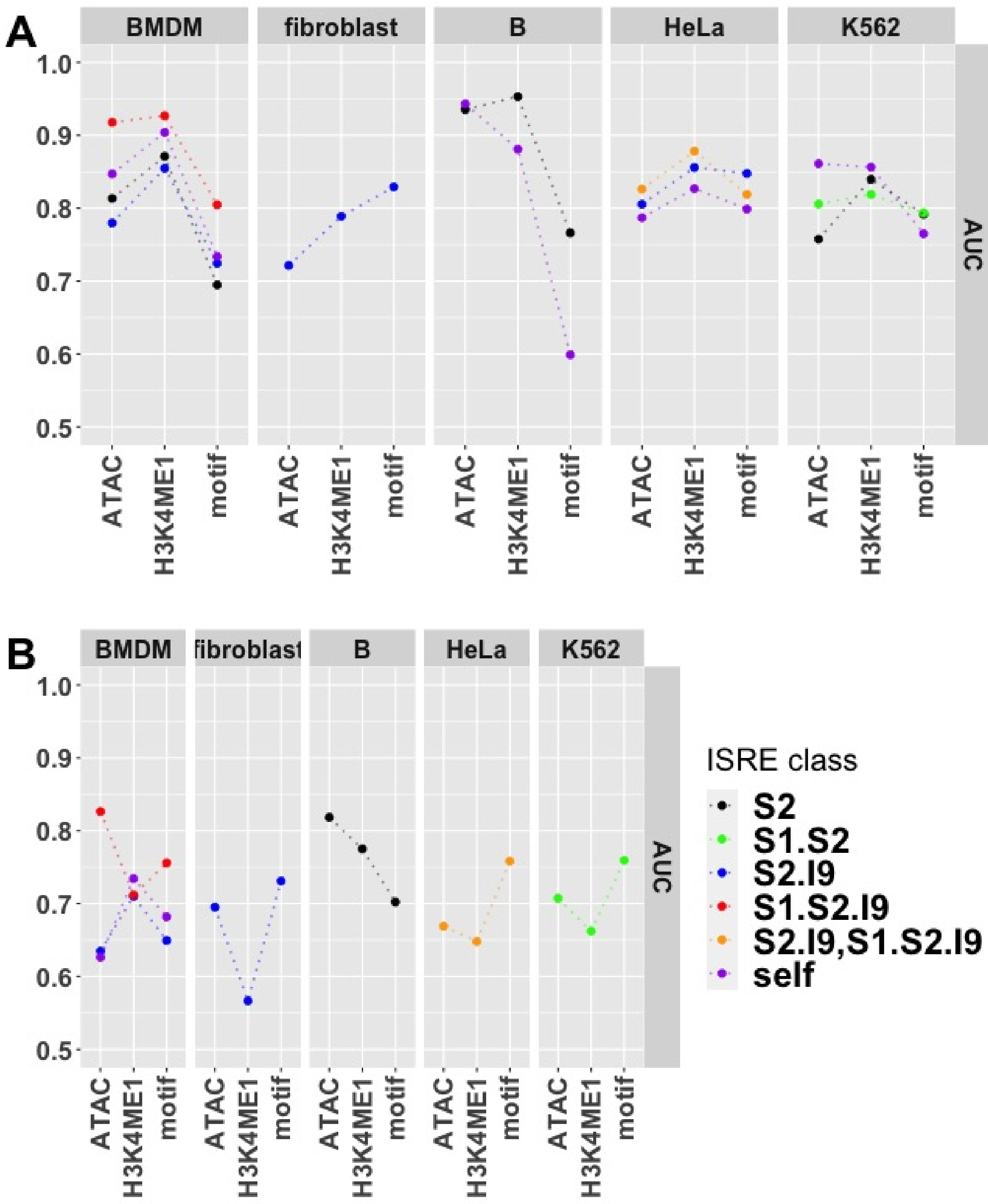


Figure S3: **Prediction Accuracy Using Transcription Factor Motifs**. We scored matches between JASPAR motifs and the 1010 base pair sequences of the ISRE loci and used the results to predict ISRE classes for ISRE in the enhancer region (A) and promoter region (B). Shown are AUC values for prediction using motifs versus using ATACseq or H3K4ME1 signals. AUC values are for prediction classifying bound ISRE against non-peak ISRE.
